# Distinct temporal expression of GW182 in neurons regulates dendritic arborization

**DOI:** 10.1101/2020.12.05.412932

**Authors:** Bharti Nawalpuri, Ravi Muddashetty

## Abstract

Precise development of the dendritic architecture is a critical determinant of mature neuronal circuitry. MicroRNA-mediated regulation of protein synthesis plays a crucial role in dendritic morphogenesis but the role of miRISC protein components in this process is less studied. Here, we show an important role of a key miRISC protein GW182 in the regulation of dendritic growth. We have identified a distinct brain region specific Spatio-temporal expression pattern of GW182 during rat postnatal development. We found that the window of peak GW182 expression coincides with the period of extensive dendritic growth, both in the hippocampus and cerebellum. Perturbation of GW182 function during a specific temporal window resulted in reduced dendritic growth of cultured hippocampal neurons. Mechanistically, we show that GW182 modulates dendritic growth by regulating global somato-dendritic translation, and actin cytoskeletal dynamics of developing neurons. Furthermore, we found that GW182 affects dendritic architecture by regulating the expression of actin modulator LIMK1. Taken together, our data reveal a previously undescribed neurodevelopmental expression pattern of GW182 and its role in dendritic morphogenesis, through both translational control and actin cytoskeletal rearrangement.

**Summary:** GW182 is a key component of miRNA induced silencing complex (miRISC). Nawalpuri et al. show that GW182 has a unique temporal expression profile during neuronal development. The developmentally controlled expression of GW182 influences dendritic morphology by regulating the expression of actin modulator LIMK1.

## Introduction

Dendrites are the primary sites of information reception and integration in neurons. Neurons from different brain regions exhibit strikingly distinct dendritic morphologies, suited to their respective functions (Hausser, 2000). Thus, precise development of the dendritic architecture is crucial for the establishment of neuronal circuitry. Consequently, defects in dendritic arborization are a common feature of multiple neurodevelopmental disorders including autism spectrum disorders (ASD) as well as intellectual disabilities (ID) (Kulkarni and Firestein, 2012; Martínez-Cerdeño, 2017).

The development of dendritic arborization is a multistep process with overlapping events of polarity establishment, dendritic extension, dendritic branching, pruning as well as stabilization (Arikkath, 2012). These events are controlled by intrinsic factors as well as extracellular cues via transcriptional and post-transcriptional gene regulatory mechanisms (Dong et al., 2015; Jan and Jan, 2010). Among these, translation regulation is one of the key mechanism employed by neurons for controlling dendritic development (Chihara et al., 2007; Jaworski et al., 2005; Kumar et al., 2005; Lein and Higgins, 1991; Ravindran et al., 2019; Slomnicki et al., 2016; Xing et al., 2012). Multiple components of the translation machinery especially ribosomes have been shown to localize to dendrites and dendritic growth cone, indicating the crucial role of local translation in dendritic development and function (Crino and Eberwine, 1996; Tiedge and Brosius, 1996). Very often, these translation regulatory pathways converge on to modulation of cytoskeletal dynamics for the regulation of dendritic arborization (Perycz et al., 2011; Ravindran et al., 2019)

Evidence highlighting the importance of translation regulation in dendritic development comes primarily from the studies involving RNA binding proteins (RBPs). Multiple translation regulatory RBPs, such as ZBP, Pumilio, Nanos, CPEB, FMRP, MOV10, and Stufen have been implicated in the regulation of dendritic development (Bestman and Cline, 2008; Brechbiel and Gavis, 2008; Lee et al., 2003; Perycz et al., 2011; Ye et al., 2004). Notably, many of these RBPs modulate the translation of cytoskeletal regulators for controlling dendritic morphogenesis. The Zipcode binding protein (ZBP) regulates dendritic branching by controlling dendritic transport and translation of β-actin (Perycz et al., 2011). FMRP is known to restrict dendritic branching in drosophila dendritic arborization (da) sensory neurons by regulation of small GTPase RAC1 mRNA to modulate dendrite branching (Lee et al., 2003). Furthermore, MOV10 mediated regulation of actin cytoskeletal proteins has been implicated in the regulation of dendritic morphogenesis (Skariah et al., 2017). Interestingly, many of these RBPs such as FMRP, Pumilio, MOV10, and CPEB act as ancillary components of miRNA induced silencing complex (miRISC) to regulate miRNA mediated gene silencing (Ford et al., 2019; Kenny et al., 2014; Nawalpuri et al., 2020; Sternburg et al., 2018).

MicroRNAs are 21-23 nucleotide long, non-coding RNA molecules involved in post-transcriptional regulation of gene expression (Bartel, 2018a; Gebert and MacRae, 2019). miRNAs are highly abundant in the nervous system and are shown to regulate the development and function of neuronal circuitry (Kosik, 2006; McNeill and Van Vactor, 2012; Rajman and Schratt, 2017; Ramakrishna and Muddashetty, 2019). The regulatory functions of microRNAs are executed by associating with a multi-protein complex called the miRISC. This complex consists of a cognate miRNA required for target mRNA recognition, along with multiple protein components to perform the gene-silencing function. GW182 and AGO2 are core miRISC proteins that work in conjunction with the ancillary components such as CCR4-NOT complex, DCP, XRN1, MOV10, and FMRP (Bartel, 2018b; Duchaine and Fabian, 2019; Filipowicz et al., 2008). These protein components associate to form functionally diverse miRISC complexes (Nawalpuri et al., 2020). The imperfect complementarity between metazoan miRNA-mRNA pairs prevents the endonucleolytic action of AGO2, causing the requirement for additional factors for effective miRNA mediated gene silencing. GW182, a large scaffolding protein, performs the essential task of recruiting mRNA deadenylation, decapping, and degradation machinery to the AGO2-miRNA-mRNA complex (Eulalio et al., 2008). This process often leads to the degradation of target mRNA (Bartel, 2018a; Gebert and MacRae, 2019). Alternatively, the AGO2-miRNA-mRNA complex can bind to RNA binding proteins such as FMRP and MOV10 leading to reversible repression of mRNA translation (Duchaine and Fabian, 2019; Gebert and MacRae, 2019; Nawalpuri et al., 2020). This suggests that miRISC composition plays a pivotal role in determining its mode of action. Interestingly, previous studies have demonstrated an important role of modulation of miRISC composition in the regulation of synaptic plasticity (Banerjee et al., 2009; Kute et al., 2019; Muddashetty et al., 2011; Rajgor et al., 2018). However, the significance of miRISC composition has not been studied in the context of neuronal development. Regulation of the expression of individual miRISC components is one of the several ways of modulating miRISC composition during neuronal development. Hence, we examined the expression pattern of several miRISC components (core as well as ancillary). In our study, we further focused on GW182 due to its central role in determining the mode of action of miRISC.

GW182, an evolutionarily conserved scaffolding protein, is a key component of metazoan miRISC (Ding and Han, 2007; Niaz and Hussain, 2018; Zielezinski and Karlowski, 2015). It is involved in the recruitment of deadenylation, decapping, and degradation enzymes to the AGO2-miRNA-mRNA complex (Eulalio et al., 2009). Structurally, it contains an N-terminal AGO binding domain rich in Glycine-tryptophan residues and a C-terminal silencing domain involved in the recruitment of mRNA degradation machinery (Eulalio et al., 2009, 2008; Pfaff et al., 2013). The C-terminal domain of GW182 also binds to the PABP complex, preventing circularization of mRNA (Huntzinger et al., 2013; Zekri et al., 2009). GW182 often localizes to cytoplasmic domains termed as Processing (P) bodies, which are non-membrane bound messenger ribonucleoprotein (mRNP) aggregates containing mRNA decay factors, translational repressors, and mRNAs (Liu et al., 2005; Luo et al., 2018). P body localization of GW182 is an important factor in driving miRNA-mediated gene silencing (Liu et al., 2005). Despite recent advances in the establishment of GW182 as a central miRISC component, the fundamental role of GW182 in neuronal development and function remains obscure. Previous studies have demonstrated the dendritic localization of GW182 (Cougot et al., 2008). Furthermore, recent studies have implicated an important role of GW182 in the regulation of BDNF induced dendritic growth in mature neurons (Huang et al., 2012). However, a comprehensive understanding of GW182 expression profile and function in neuronal development is lacking.

In the current study, we describe a novel function of miRISC protein GW182 in the regulation of dendritic arborization in the hippocampus. Our analysis of miRISC components expression profile revealed a distinct brain-specific Spatio-temporal expression pattern of GW182. Our results indicate that GW182 regulates dendritic growth by controlling actin dynamics through the regulation of LIMK1 expression.

## Results

### GW182 expression follows a distinct Spatio-temporal profile during neuronal development

We investigated the expression pattern of miRISC components AGO2, GW182, XRN1, MOV10, and FMRP during rat brain development in two distinct brain regions, hippocampus and cerebellum. The timeline of different neurodevelopmental processes is well characterized in both these regions, providing the opportunity for correlating the changes in the expression pattern of miRISC proteins to specific neurodevelopmental stages such as dendritogenesis and synapse morphogenesis. Notably, maturation of the cerebellar circuitry is protracted as compared to the hippocampal region (excluding dentate gyrus).

Rat hippocampal and cerebellar tissues were harvested at different stages of brain development starting from embryonic day 18 (E18) to postnatal day 60 (P60) and were subjected to immunoblotting to analyze the expression of miRISC protein components **(Figure 1A)**. In the hippocampus, the levels of AGO2 significantly increased during postnatal development from E18 to P60 **(Figure 1B)**, while the expression of MOV10 and XRN1 remained unchanged **(Figure S1B, S1C)**. FMRP expression increased 2.5 times from P5 to P15 and remained unchanged from P15 onwards **(Figure S1A)**. Interestingly, hippocampal expression of GW182 peaked at P7, followed by a 10-fold reduction in expression from P7 to P15. This reduced expression was sustained from P15 onwards **(Figure 1C)**.

**Figure 1:**
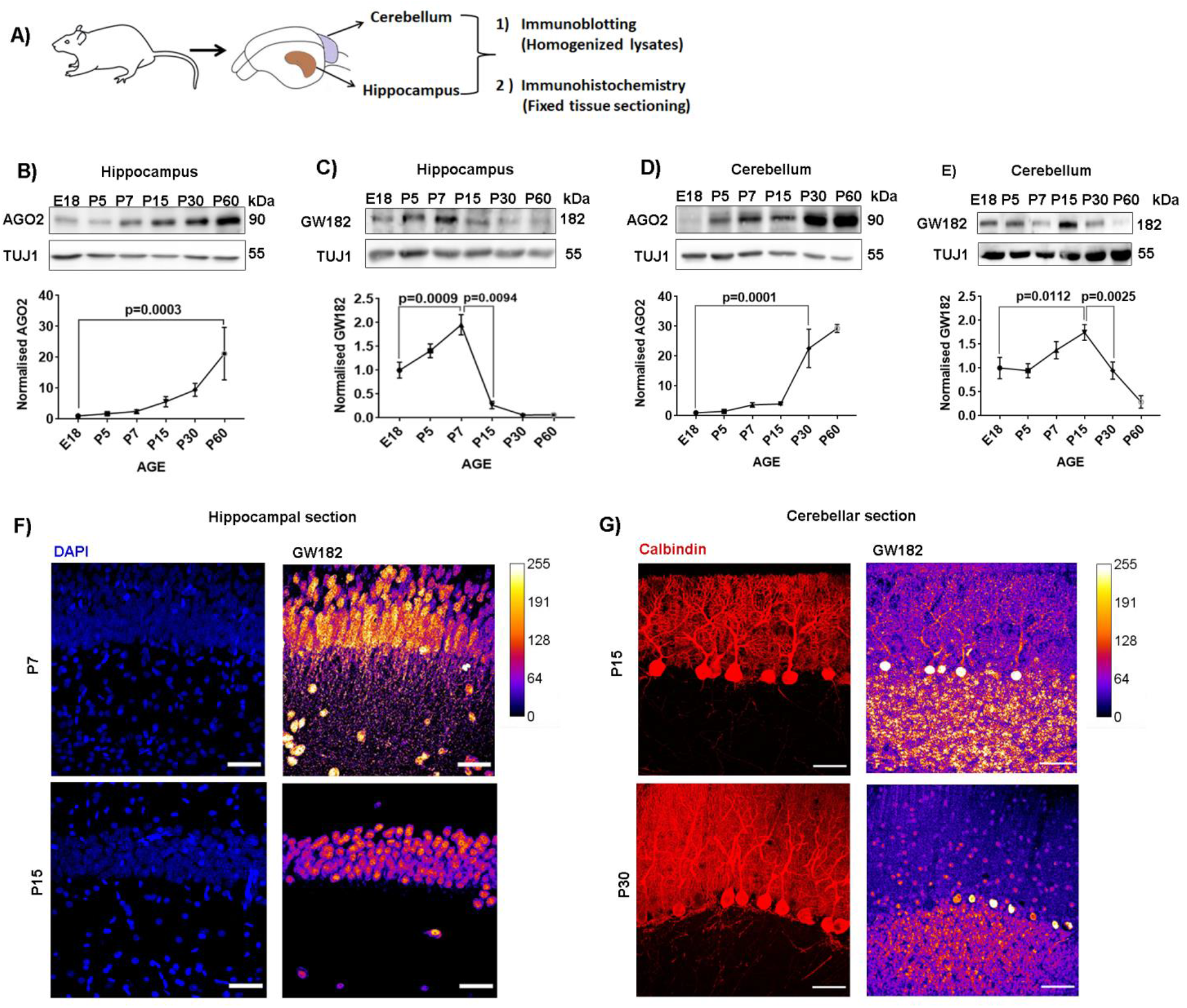
GW182 expression is developmentally regulated in rodent brain. A) Schematic showing the experimental procedure for assessment of miRISC proteins expression pattern from rodent brain. B) Representative immunoblots (top) and line graph (bottom) depicting AGO2 expression profile during hippocampal development. Data represent relative AGO2 levels normalized to TUJ1. Data: mean +/- SEM, n=3-5 animals per group, One Way ANOVA followed by Bonferroni’s multiple comparisons test. C) Representative immunoblots (top) and line graph (bottom) depicting GW182 expression profile during hippocampal development. Data represent relative GW182 levels normalized to TUJ1. Data: mean +/- SEM, n=3-5 animals per group, One Way ANOVA followed by Bonferroni’s multiple comparisons test. D) Representative immunoblots (top) and line graph (bottom) depicting AGO2 expression profile during Cerebellar development. Data represent relative AGO2 levels normalized to TUJ1. Data: mean +/- SEM, n=3-5 animals per group, One Way ANOVA followed by Bonferroni’s multiple comparisons test. E) Representative immunoblots (top) and line graph (bottom) depicting GW182 expression profile during Cerebellar development. Data represent relative GW182 levels normalized to TUJ1. Data: mean +/- SEM, n=3-5 animals per group, One Way ANOVA followed by Bonferroni’s multiple comparisons test. F) Representative immunohistochemistry images showing DAPI and GW182 staining in rat hippocampal CA1 region at P7 and P15. Scale bar represents 50 microns. G) Representative immunohistochemistry images showing Calbindin and GW182 staining in rat Cerebellar sections at P15 and P30. Scale bar represents 50 microns.

With cerebellar lysates, similar to the hippocampus, we found that AGO2 expression gradually increased from E18 to P60 **(Figure 1D)**. Similarly, XRN1 expression also increased significantly from P5 to P30 **(Figure S1F)**. The expression of MOV10 peaked at P7, with a sustained reduction from P15 onward **(Figure S1E)**. Similar to the hippocampus, FMRP expression also increased 2.5 times from P5 to P15, however, the levels reduced at P30 **(Figure S1D)**. In the cerebellum, GW182 levels peaked at P15, with a gradual reduction from P15 to P60 **(Figure 1E)**. Careful introspection of immunoblotting results from both cerebellum and hippocampus highlights one striking feature; in both the regions, the expression of GW182 peaks during the time of extensive dendritic arborization, followed by a significant reduction of expression in the mature tissue. Among all the proteins examined only GW182 showed this peculiar expression profile. These findings suggest an important role of GW182 in neuronal development.

To understand if the distinct expression pattern of GW182 is a brain-specific feature, we investigated the expression profile of GW182 in the liver lysate. In the liver, GW182 levels were almost undetectable at P5, with a significant increase from P5 to P15 **(Figure S1G)**. The levels remained elevated thereafter. This finding supports the idea of the brain-specific temporal profile of GW182 expression and function. We also investigated the developmental expression profile of GW182 mRNA, to examine if it mirrors the protein expression pattern. In the hippocampus, the levels of GW182 mRNA remained unchanged between P7 to P15 (Point of peak GW182 expression and period of sustained drop) **(Figure S1I)** In contrast, the AGO2 mRNA expression profile mirrored the protein expression pattern of AGO2. **(Figure S1H)**. A similar trend was observed in the cerebellum as well, where GW182 mRNA levels remained unchanged from P15 to P60 **(Figure S1K)**, whereas AGO2 mRNA levels increased significantly at the time point examined **(Figure S1J)**.

We further investigated the spatial localization (somato-dendritic) of GW182 during neuronal development. For this, we performed immunohistochemistry in the hippocampal CA1 region at P7 and P15 (Point of peak GW182 expression and period of sustained drop in the hippocampus). The primary cell body layer *stratum pyramidale* was identified with DAPI staining. At P7, GW182 staining was observed in both *stratum pyramidale* (cell body) as well as in *stratum oriens* (dendritic layer) **(Figure 1F)**. At P15, we observed reduced GW182 staining in both the layers, with a pronounced reduction in stratum oriens **(Figure 1F)**. We also examined GW182 expression in cerebellar slices at P15 and P30 (Point of peak GW182 expression and period of drop in the cerebellum). Calbindin was used as a marker to identify Purkinje neuron cell bodies and dendrites. At P15, we observed GW182 expression in all three layers of cerebellum, namely granular layer, Purkinje layer, and molecular layer **(Figure 1G)**. On the contrary, at P60, we observed a drastic reduction in GW182 levels, which was most pronounced in the molecular layer **(Figure 1G)**.

Altogether, these results demonstrate the distinct, brain-specific Spatio-temporal expression profile of miRISC protein GW182. The levels of GW182 peaked during the period of extensive dendritogenesis, followed by a reduction in GW182 levels in adult neuronal circuitry.

### Spatio-temporal expression pattern of GW182 is recapitulated in hippocampal neuronal culture

In the previous experiments, we demonstrated that GW182 follows a distinct Spatio-temporal expression profile during brain development. Since GW182 levels peaked during the dendritogenesis period, we wanted to study the role of GW182 in the regulation of dendrite morphology. For this, we selected dissociated hippocampal neuronal cultures as our model system. This culture paradigm offers several advantages such as accessibility and amenability to molecular and pharmacological manipulations. We first investigated if the Spatio-temporal expression pattern of GW182 is recapitulated in a dissociated culture system. Hippocampal neurons were cultured from embryonic day 18 (E18) rat embryos as previously described (Kaech and Banker, 2006) **(Figure 2A)**. Cultured neurons were harvested at different developmental stages (Days in vitro (DIV) 3, 6, 9, 12, 15, and 21) and subjected to immunoblotting analysis for GW182 protein.

**Figure 2:**
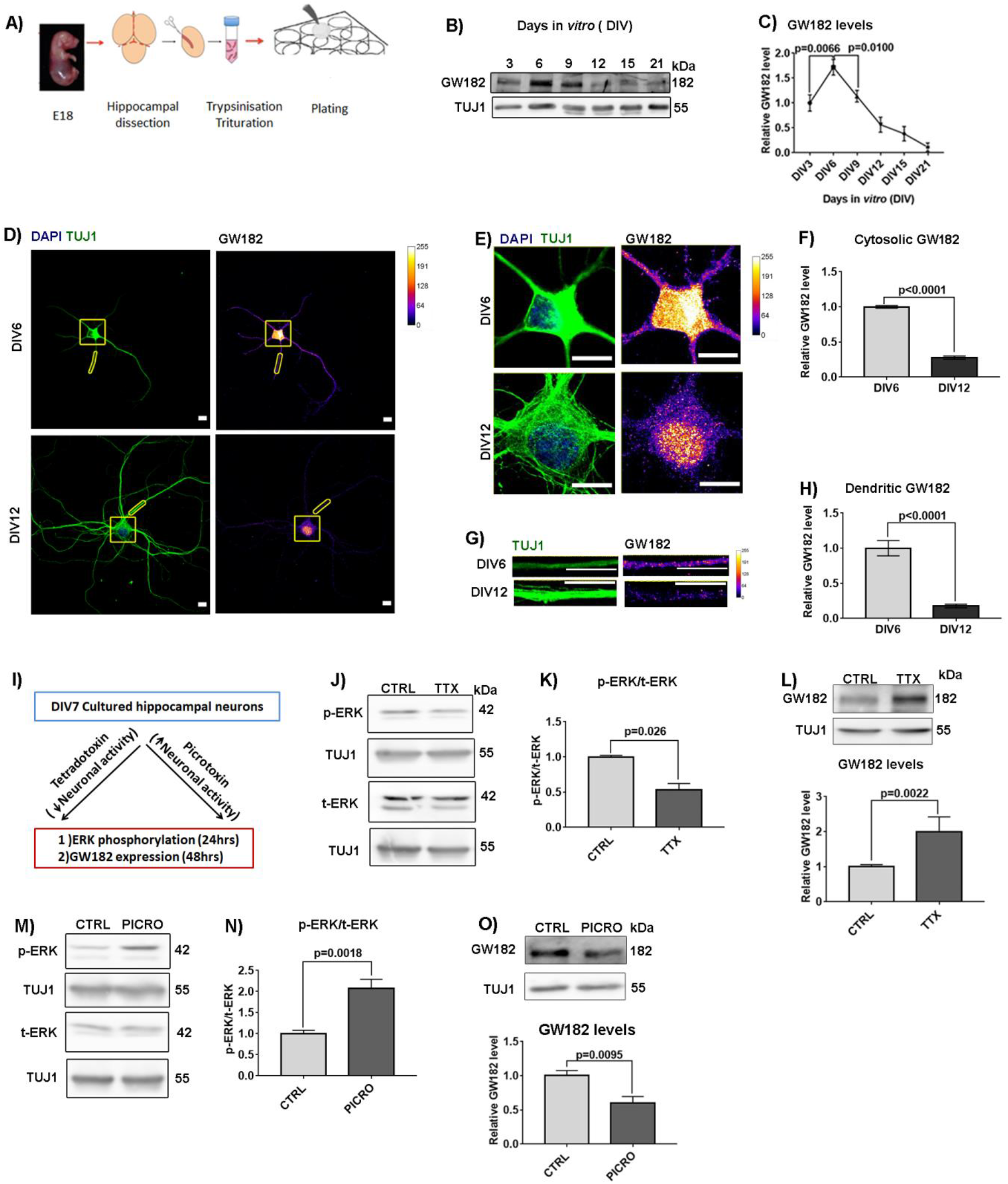
Spatio-temporal expression pattern of GW182 in culture hippocampal neurons. A) Schematic representing the procedure for embryonic hippocampal neuronal culture. B) Representative immunoblots depicting GW182 expression profile in hippocampal neuronal culture from DIV3 to DIV21. C) Line graph depicting GW182 expression profile in hippocampal neuronal culture. Data represent relative GW182 levels normalized to TUJ1. Data: mean +/- SEM, n=4-6 independent experiments, One Way ANOVA followed by Bonferroni’s multiple comparisons test. D) Representative image showing DAPI, TUJ1, and GW182 immunostaining in cultured neurons at DIV 6 and DIV12. E) Enlarged image of a section of cell body from Figure 2D showing DAPI, TUJ1, and GW182 immunostaining in cultured neurons at DIV 6 and DIV12. Scale bar represents 10 microns. F) Quantification of normalized mean intensity of cytosolic GW182 levels in DIV6 and DIV12 neurons, Data: mean +/- SEM n=27-32 neurons from 4 independent experiments, Mann-Whitney test. G) Enlarged image of a dendritic segment from Figure 2D showing DAPI, TUJ1, and GW182 immunostaining in cultured neurons at DIV 6 and DIV12. Scale bar represents 10 microns. H) Quantification of normalized mean intensity of dendritic GW182 levels in DIV6 and DIV12 hippocampal neurons, Data: mean +/- SEM n=27-32 neurons from 4 independent experiments, Mann-Whitney test. I) Schematic representing the picrotoxin or tetradotoxin treatment protocol in cultured hippocampal neurons. J) Representative immunoblots depicting ERK phosphorylation status 24hrs post tetradotoxin treatment of DIV7 hippocampal neuronal culture. K) Quantification of ERK phosphorylation after 24hrs tetradotoxin treatment of DIV7 hippocampal neuronal culture. Data represent the ratio of phospho/total (p/t) ERK levels normalized to TUJ1. Data: mean +/- SEM, n=5 independent experiments, Unpaired t-test. L) Representative immunoblots (top) and corresponding quantification (bottom) depicting GW182 levels 48hrs post tetradotoxin treatment in hippocampal neuronal culture. Data represent relative GW182 levels normalized to TUJ1. Data: mean +/- SEM, n=5 independent experiments, Unpaired t-test. M) Representative immunoblots depicting ERK phosphorylation post 24hr Picrotoxin treatment of DIV7 hippocampal neuronal culture. N) Quantification of ERK phosphorylation on 24hr picrotoxin treatment of DIV7 hippocampal neuronal culture. Data represent the ratio of relative p/T ERK levels normalized to TUJ1. Data: mean +/- SEM, n=5 independent experiments, Unpaired t-test. O) Representative immunoblots (top) and corresponding quantification (bottom) depicting GW182 levels post 48hr picrotoxin treatment in hippocampal neuronal culture. Data represent relative GW182 levels normalized to TUJ1. Data: mean +/- SEM, n=5 independent experiments, Unpaired t-test.

GW182 expression could be detected in cultured neurons as early as DIV3. The levels peaked at DIV6 followed by a drastic reduction at DIV9. GW182 levels further reduced significantly from DIV9 to DIV12 **(Figure 2B, 2C)**. The reduced expression of GW182 was maintained in DIV21 mature neurons. Thus, the temporal expression of GW182 in cultured neurons is parallel to its in vivo expression. Next, we investigated the spatial expression pattern of GW182 in cultured neurons using quantitative immunostaining **(Figure 2C)**. We measured cytosolic and dendritic levels of GW182 in cultured neurons at DIV6 (point of peak expression) and DIV 12 (period after which GW182 levels do not drop further). We observed a significant reduction in cytosolic as well as dendritic levels of GW182 from DIV6 to DIV12 **(Figure 2D, 2E, 2F, and 2G, 2H)**. Hereby, we conclude that the Spatio-temporal expression pattern of GW182 is recapitulated in dissociated hippocampal neuronal cultures. Hence, neuronal cultures can be used to study the functional role of GW182.

Previously, we observed that GW182 expression peaks during the period of extensive dendrite growth, and its level begin to drop when the neurons become primed to respond to neuronal activity (beginning of synaptogenesis). We hypothesized that the increase in neuronal activity during this developmental period drives the reduction in GW182 expression. To test this, DIV7 neuronal cultures were chronically treated either with a GABA_(A)_ receptor antagonist picrotoxin to increases neuronal activity, or with sodium channel blocker tetradotoxin (TTX) for suppression of neuronal activity **(Figure 2I)**. We validated tetradotoxin treatment by quantifying levels of ERK phosphorylation. In accordance with previous reports (Bateup et al., 2013), we observed over 50% reduction in ERK phosphorylation on 24hr of tetradotoxin treatment **(Figure 2J and 2K)**. When we tested the effect of tetradotoxin treatment on GW182 levels, we found that 48 hrs of tetradotoxin treatment results in an over two-fold increase in GW182 protein levels **(Figure 2L)**. GW182 mRNA also showed a similar trend of increase (although not significant) in response to tetradotoxin treatment **(Figure S2A)**.

Conversely, increased ERK phosphorylation (24hrs) was used to validate picrotoxin treatment **(Figure 2M, 2N, and S2A)**. We observed around 40% reduction in GW182 levels upon 48hrs of Picrotoxin treatment **(Figure 2O)**. The reduced protein expression on picrotoxin treatment was captured at the mRNA level as well **(Figure S2B)**. In summary, we found that GW182 levels are inversely regulated by neuronal activity.

Altogether, these results establish the recapitulation of the GW182 expression pattern in dissociated neuronal cultures. Furthermore, we verify the role of neuronal activity in the regulation of GW182 expression.

### GW182 loss of function reduces the complexity of dendritic arbors

The developmentally controlled expression and dendritic localization of GW182 indicate its possible role in the regulation of dendritic arborization. To address this, we utilized a loss of function approach. We used a GFP-tagged dominant negative mutant of GW182 to perturb its function (Jakymiw et al., 2005) during a period of rapid dendritogenesis (DIV3-7) in cultured neurons. GW182 is a large scaffolding protein that consists of an N-terminal AGO binding domain and a C-terminal silencing domain which performs the effector function of GW182. The dominant negative mutant lacks the silencing domain and hence cannot function in the canonical miRNA pathway (Jakymiw et al., 2005).

The expression of GFP-tagged dominant negative (DN) GW182 was verified in HEK 293T cells using immunoblotting and immunostaining analysis **(Figure S5A and S5C)**. The dominant negative mutant gives a band of around 85 kilodaltons (kDa) in western blotting on probing with GFP antibody **(Figure S5A)**. Immunostaining analysis revealed a diffused expression pattern of DNGW182 in contrast to full-length GW182 (overexpressed) which shows punctate distribution **(Figure 5SC)**. After initial validation, we overexpressed GFP DNGW182 in primary hippocampal neurons on DIV3 and assessed its effect on dendritic arborization on DIV7 **(Figure 3A)**. Neurons were selected based on GFP staining, and dendrites were identified with MAP2 immunostaining. Neurons overexpressing GFP construct (without GW182) were used as control. We used Sholl analysis for studying dendrite arborization and Neuron J neurite tracing software for quantifying the dendrite length. Neurons overexpressing DNGW182 showed a downward shift in Sholl profile as compared to GFP transfected neurons, indicating a less complex dendritic arbor **(Figure 3B and 3C)**. These neurons also showed significantly reduced dendritic intersections at a distance of 8-64 microns from the soma. We also found a reduced number of total dendritic intersections as well as a reduction in total dendritic length in neurons overexpressing DNGW182 **(Figure S3A and 3D)**. In summary, perturbation of GW182 function using DNGW182 resulted in reduced complexity of dendritic arborization.

**Figure 3:**
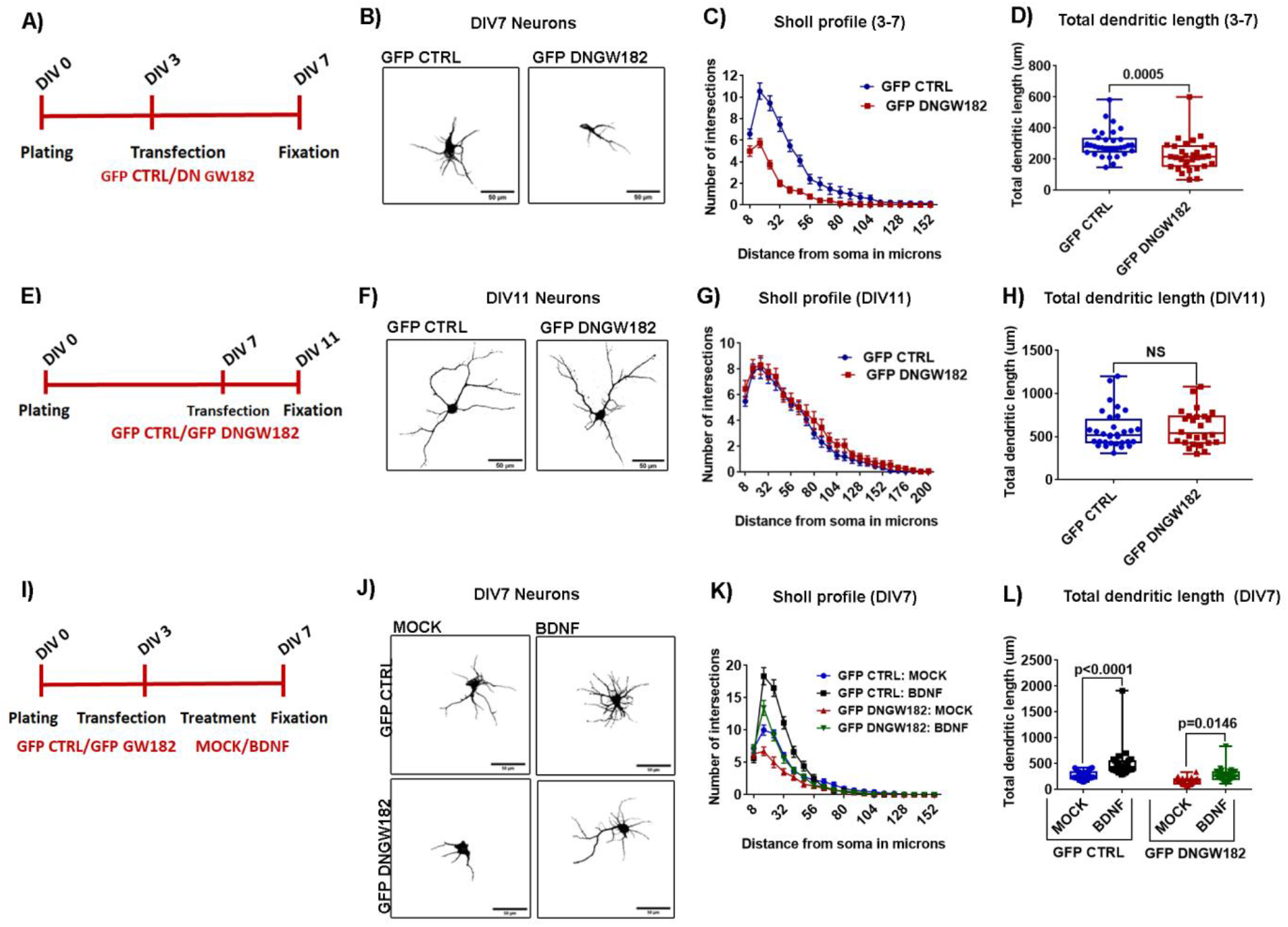
Overexpression of DNGW182 mutant decreases dendritic arborization during a distinct neurodevelopmental window. A) Schematic showing the experimental procedure and timeline: Cultured hippocampal neurons were transfected with either control GFP or GFP DNGW182 on DIV3 and fixed on DIV7, followed by immunostaining. Dendrites were identified with MAP2 staining. B) Representative micrographs of DIV7 cultured hippocampal neurons transfected with either control GFP or GFP DNGW182. The images are derived from the threshold MAP2 intensity of transfected neurons. Scale bar represents 50 microns. C) Intersection profile / Sholl curve of DIV7 cultured hippocampal neurons transfected with either control GFP or GFP DNGW182. Data: mean +/- SEM, n=32 neurons from 4 independent experiments, GFP DNGW182 overexpressing neurons had significantly more dendrites from GFP overexpressing neurons at 16-48 microns from the soma, Two way ANOVA followed by Bonferroni’s multiple comparisons test. D) Quantification of the total dendritic length of cultured hippocampal neurons transfected with either control GFP vector or DN GW182, n=27-32 neurons from 4 independent cultures, Mann-Whitney test. E) Schematic showing the experimental procedure and timeline: Cultured hippocampal neurons were transfected with either control GFP or GFP DNGW182 on DIV7 and fixed on DIV11, followed by immunostaining. Dendrites were identified with MAP2 staining F) Representative micrographs of DIV11 cultured hippocampal neurons transfected with either control GFP vector or GFP DNGW182 mutant at DIV7. The images are derived from the threshold MAP2 intensity of transfected neurons. Scale bar represents 50 microns. G) Intersection profile / Sholl curve of DIV11 cultured hippocampal neurons transfected with either control GFP or GFP DNGW182 mutant. Data: mean +/- SEM, n=22-25 neurons from 4 independent experiments. H) Quantification of the total dendritic length of DIV11 cultured hippocampal neurons transfected with either Control GFP or GFP DNGW182, n=22-23 neurons from 4 independent cultures, Unpaired t-test with Welch’s correction. I) Schematic showing the experimental procedure and timeline: Cultured hippocampal neurons were transfected on DIV3 followed by BDNF treatment for 48hrs starting from DIV5 onwards, and fixed on DIV7. J) Representative images of cultured hippocampal neurons transfected at DIV3 with either control GFP or GFP DNGW182. After transfection, the neurons were treated with BDNF (50ng/ml) on DIV 5 and were fixed on DIV7. Scale bar represents 50 microns. K) Sholl Quantification of the effects of GFP DNGW182 on BDNF-induced dendrite arborization formation of hippocampal neurons. Data: mean +/- SEM, n=25-30 neurons from 4 independent experiments, BDNF treatment resulted in increased dendritic intersections in GFP transfected neurons at 16-40 microns from the soma and resulted in increased intersections in GFP DNGW182 transfected neurons as well at 16-32 microns from the soma, Two way ANOVA followed by Bonferroni’s multiple comparisons test. L) Quantification of the total dendritic length of neurons described in Figure 3E, n=25-30 neurons from 3 independent experiments, One Way Anova followed by Bonferroni’s multiple comparisons test.

Next, we questioned if GW182 regulates dendritic morphology during later stages of dendritic development as well. To this end, we transfected DIV7 hippocampal neurons with GFP or GFP tagged DNGW182 and fixed the neurons on DIV11 for evaluation of dendritic morphology (period of rapid dendritic growth and initiation of synaptogenesis) **(Figure 3E)**. We found that at this stage, overexpression of DNGW182 did not cause any change in dendritic morphology as depicted by the Sholl profile **(Figure 3F and 3G)**. Furthermore, we did not observe any difference in the total number of dendritic intersections and total dendritic length **(Figure S3B and 3H)**. These results suggest that GW182 regulates dendritic growth only during a restricted temporal window.

Multiple studies have demonstrated the importance of BDNF in the regulation of dendrite arborization (Gorski et al., 2003; McAllister et al., 1996). Here we examined whether perturbation of GW182 function affects BDNF induced dendritic growth and arborization in cultured hippocampal neurons. Neurons were transfected with either GFP or GFP-tagged DNGW182 on DIV3 followed by 48hrs BDNF (50ng/ml) treatment from DIV5 to DIV7. The neurons were fixed on DIV7, followed by immunostaining **(Figure 3I)**. In GFP transfected neurons, BDNF stimulation resulted in enhanced dendrite arborization as demonstrated by the Sholl profile **(Figure 3J and 3K)**. Further, BDNF treatment resulted in an increase of total dendritic length and total intersections in GFP transfected neurons **(Figure 3L and S3C)**. This validated the biological response of BDNF treatment in our system. When BDNF response was measured in DNGW182 transfected neurons, we found these neurons were still responsive to BDNF stimulation **(Figure 3J, 3K, 3L, and S3C)**. Similar to GFP transfected neurons, BDNF treatment increased the dendritic arborization and length of DNGW182 transfected neurons as well **(Figure 3J, 3K, and 3L)**. BDNF stimulation restored the dendritic morphology DNGW182 overexpressing neurons to that of control neurons. These results indicated that GW182 does not modulate BDNF induced dendritic growth in DIV3-7 neurons.

As an independent loss of function approach, we used siRNA against GW182 to demonstrate its role in the regulation of dendritic morphology. The siRNA mediated knockdown of GW182 was validated with immunoblotting in Neuro2A cells. GW182 siRNA transfected cells showed around 70 percent reduction in GW182 protein levels as compared to scrambled siRNA transfected cells **(Figure S4A)**. We also validated GW182 knockdown in cultured hippocampal neurons with immunolabelling technique **(Figure S4B)**. GW182 siRNA transfection resulted in a reduction of GW182 protein as measured by immunofluorescence, with a substantial depletion of GW182 from dendritic processes but a relative sparing of somatic GW182 staining, suggesting that different pools of GW182 might have different turnover rates. To study the effect of GW182 knockdown on dendrite arborization, cultured hippocampal neurons were transfected with GFP along with scrambled siRNA or GW182 siRNA on DIV3, followed by fixation and immunostaining on DIV7 **(Figure 4A)**. Similar to the phenotype observed with DNGW182, we found that siRNA-mediated knockdown of GW182 also resulted in a reduced number of dendritic intersections in the Sholl curve, indicating a reduction in dendrite arborization **(Figure 4B and 4C)**. Moreover, neurons transfected with GW182 siRNA showed approximately 20 percent reduction in total dendritic length and total number of intersections as compared to the scrambled siRNA transfected neurons **(Figure 4D, and S4C)**. We also addressed the effect of GW182 knockdown in cultured neurons during later stages of dendritic development (DIV7-11) **(Figure 4E)**. At this stage, we did not observe any difference in the dendritic morphology between GW182 siRNA and scrambled siRNA transfected neurons **(Figure 4F and 4G)**. The neurons showed a similar number of dendritic intersections as well as total dendritic length **(Figure S4E and 4H)**. Taken together, these results showed that the loss of function of GW182 results in reduced dendritic arborization and dendritic length during a restricted temporal window of dendritic development.

**Figure 4:**
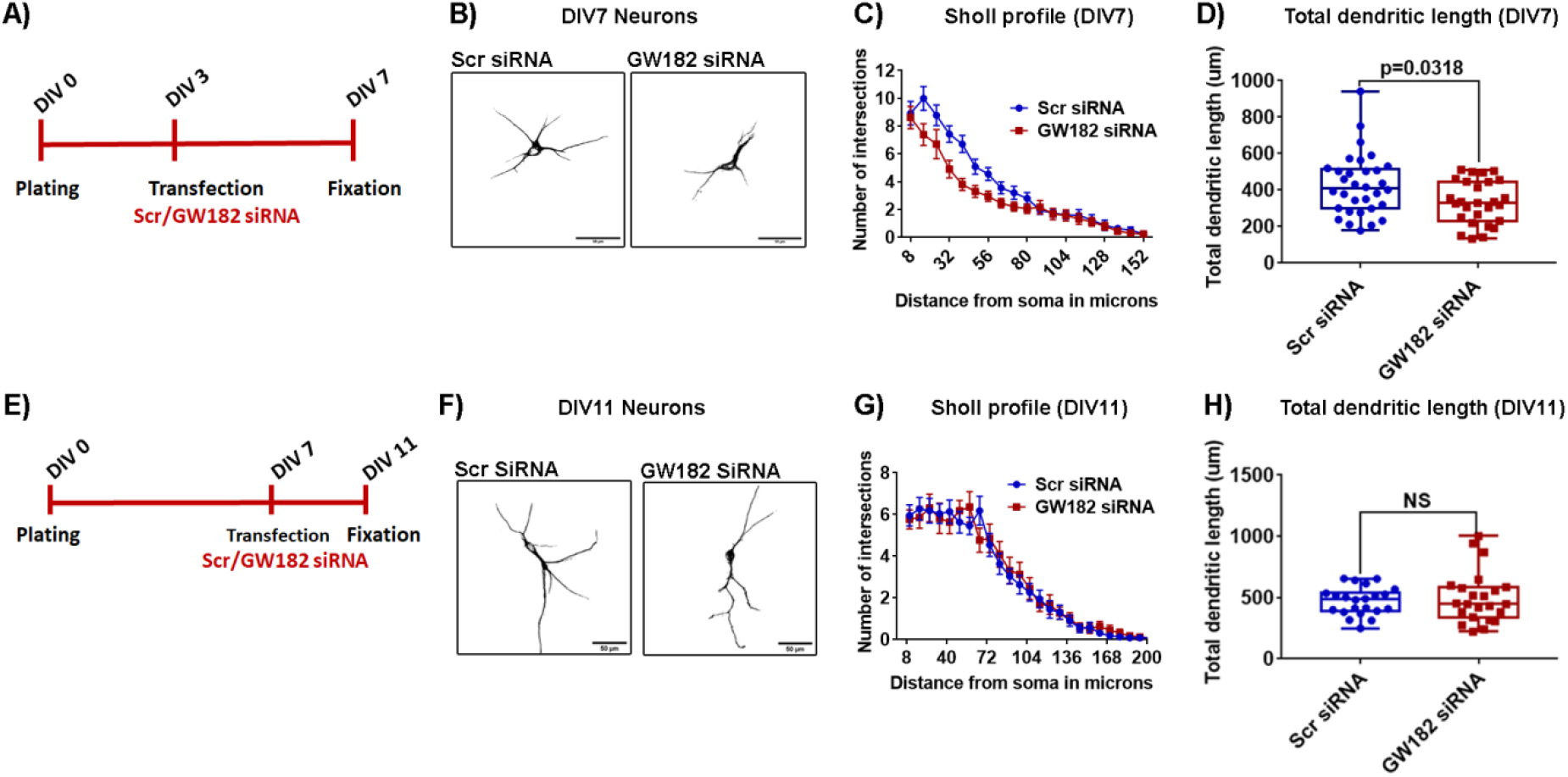
GW182 knockdown leads to a reduction in dendritic arborization of hippocampal neurons. A) Schematic showing the experimental procedure and timeline: Cultured hippocampal neurons were transfected with either scrambled siRNA or GW182 siRNA along with GFP on DIV3 and were fixed on DIV7, followed by immunostaining. Dendrites were identified with MAP2 staining (Top schematic). B) Representative micrographs of DIV7 cultured hippocampal neurons transfected with either scrambled siRNA or GW182 siRNA. The images are derived from the threshold MAP2 intensity of transfected neurons. Scale bar represents 50 microns. C) Intersection profile / Sholl curve of DIV7 cultured hippocampal neurons transfected with either scrambled siRNA or GW182 siRNA on DIV3. Data: mean +/- SEM, n=32 neurons from 4 independent experiments, GW182 siRNA overexpressing neurons had significantly more dendrites than scrambled siRNA overexpressing neurons at 16-40 microns from the soma, two way ANOVA followed by Bonferroni’s multiple comparisons test. D) Quantification of the total dendritic length of DIV7 hippocampal neurons transfected with either scrambled siRNA or GW182 siRNA on DIV3, n=27-32 neurons from 4 independent cultures, Mann-Whitney test. E) Schematic showing the experimental procedure and timeline: Cultured hippocampal neurons were transfected with either scrambled siRNA or GW182 siRNA along with GFP on DIV7 and were fixed on DIV11 (Bottom schematic). F) Representative micrographs of DIV11 hippocampal neurons transfected with either scrambled siRNA or GW182 siRNA on DIV7. The images are derived from the threshold MAP2 intensity of transfected neurons. Scale bar represents 50 microns. G) Intersection profile / Sholl curve of DIV11 cultured hippocampal neurons transfected with either Scrambled siRNA or GW182 siRNA on DIV7. Data: mean +/- SEM, n=32 neurons from 4 independent experiments. H) Quantification of the total dendritic length of DIV11 hippocampal neurons transfected with either scrambled siRNA or GW182 siRNA on DIV7, n=22-23 neurons from 4 independent cultures, Unpaired t-test with Welch’s correction.

### GW182 overexpression potentiates dendritic growth and arborization

In the above section, we used a loss of function method to demonstrate the role of GW182 in the regulation of dendritic morphology. To further strengthen our finding, we employed a gain of function approach involving overexpression of full-length GW182. The over-expression of GFP tagged GW182 was validated in HEK 293T cells **(Figure S5B and S5C).** To study the effect of GW182 overexpression on dendritic arborization, we transfected cultured hippocampal neurons on DIV3 with either GFP or GFP GW182, followed by fixation and immunostaining at DIV7 **(Figure 5A)**. Sholl analyses indicated an increase in dendritic branching in GW182 overexpressing neurons as compared to GFP transfected neurons **(Figure 5B and 5C)**. GW182 overexpression resulted in a 2 fold increase in total dendritic length as well as the total number of dendritic intersections **(Figure 5D and S5B)**. Hence, overexpression of GW182 promotes dendritic growth arborization and has an opposite effect on dendritic arborization compared to GW182 knockdown. Collectively, these experiments demonstrate the permissive role of GW182 in the regulation of dendrite morphology.

**Figure 5:**
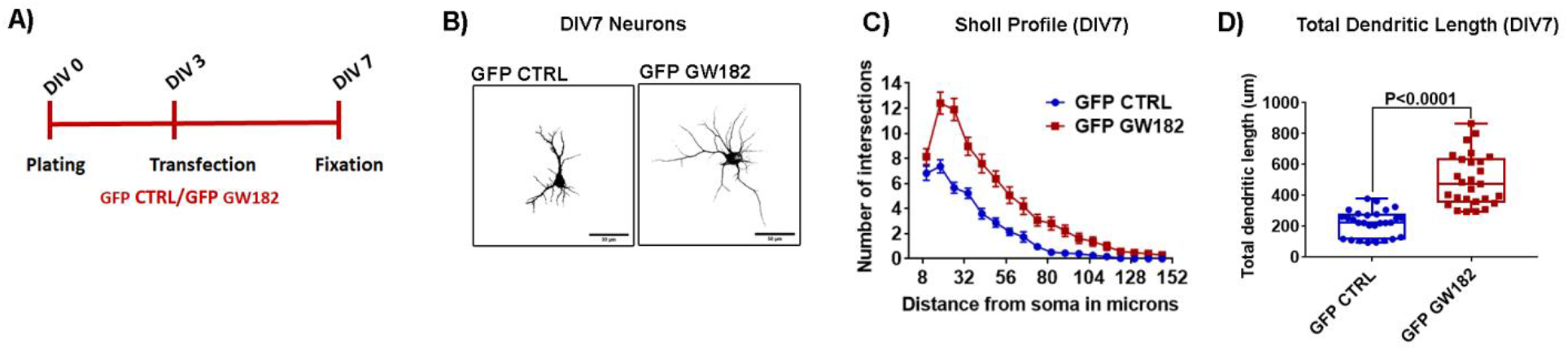
GW182 overexpression enhances dendrite growth and arborization of Hippocampal neurons. A. Schematic showing the experimental procedure and timeline: Cultured Hippocampal neurons were transfected with GFP or GFP GW182 at DIV3 and fixed at DIV7, followed by immunostaining. Dendrites were identified with MAP2 staining. B. Representative micrographs of DIV7 hippocampal neurons transfected with GFP or GFP GW182. The images are derived from the threshold MAP2 intensity of transfected neurons. Scale bar represents 50 microns. C. Intersection profile / Sholl curve of DIV7 neurons transfected with GFP or GFP GW182. Data: mean +/- SEM, n=32 neurons from 4 independent experiments, GFP DNGW182 overexpressing neurons had significantly more dendrites from GFP overexpressing neurons at 16-80 microns from the soma, two way ANOVA followed by Bonferroni’s multiple comparisons test. D. Quantification of the total dendritic length of DIV7 hippocampal neurons transfected with either GFP or GFP GW182 vector. n=27 neurons from 3 independent cultures, Mann-Whitney test.

### GW182 regulates somatodendritic translation in developing neurons

We next sought to identify the molecular mechanism of GW182 mediated regulation of dendritic growth. Several studies have demonstrated the importance of protein synthesis and various RNA binding proteins in the regulation of diverse aspects of dendrite morphogenesis (Bestman and Cline, 2008; Chihara et al., 2007; Perycz et al., 2011; Ye et al., 2004). Furthermore, multiple studies have also implicated an important role of GW182 in translational silencing (Yao et al., 2011). Based on the above information, we wanted to investigate the role of GW182 in the regulation of neuronal translation. In our previous experiments, we demonstrated that the alterations in GW182 expression at DIV3 results in dendritic morphology changes at DIV7. We thereby reasoned that GW182 might regulate neuronal translation at an earlier time point to influence dendrite morphology at DIV7. Therefore, we transfected the neurons with either GW182 siRNA/scrambled siRNA for GW182 knockdown, or overexpressed GFP/GFP GW182/GFP DN GW182 at DIV3, followed by the assessment of neuronal translation at DIV5/DIV6 **(Figure 6A)**. We monitored neuronal translation using FUNCAT metabolic labeling technique as described previously (Dieck et al., 2012; Dieterich et al., 2010). The extent of labeling was quantified using mean fluorescence intensity which proportionally correlates with the rate of protein synthesis throughout the labeling period. The reduction in the FUNCAT signal of neurons incubated with the protein synthesis inhibitor anisomycin verified that the signal is specifically attributed to the newly synthesized proteins during the labeling period **(Figure S6A)**.

**Figure 6:**
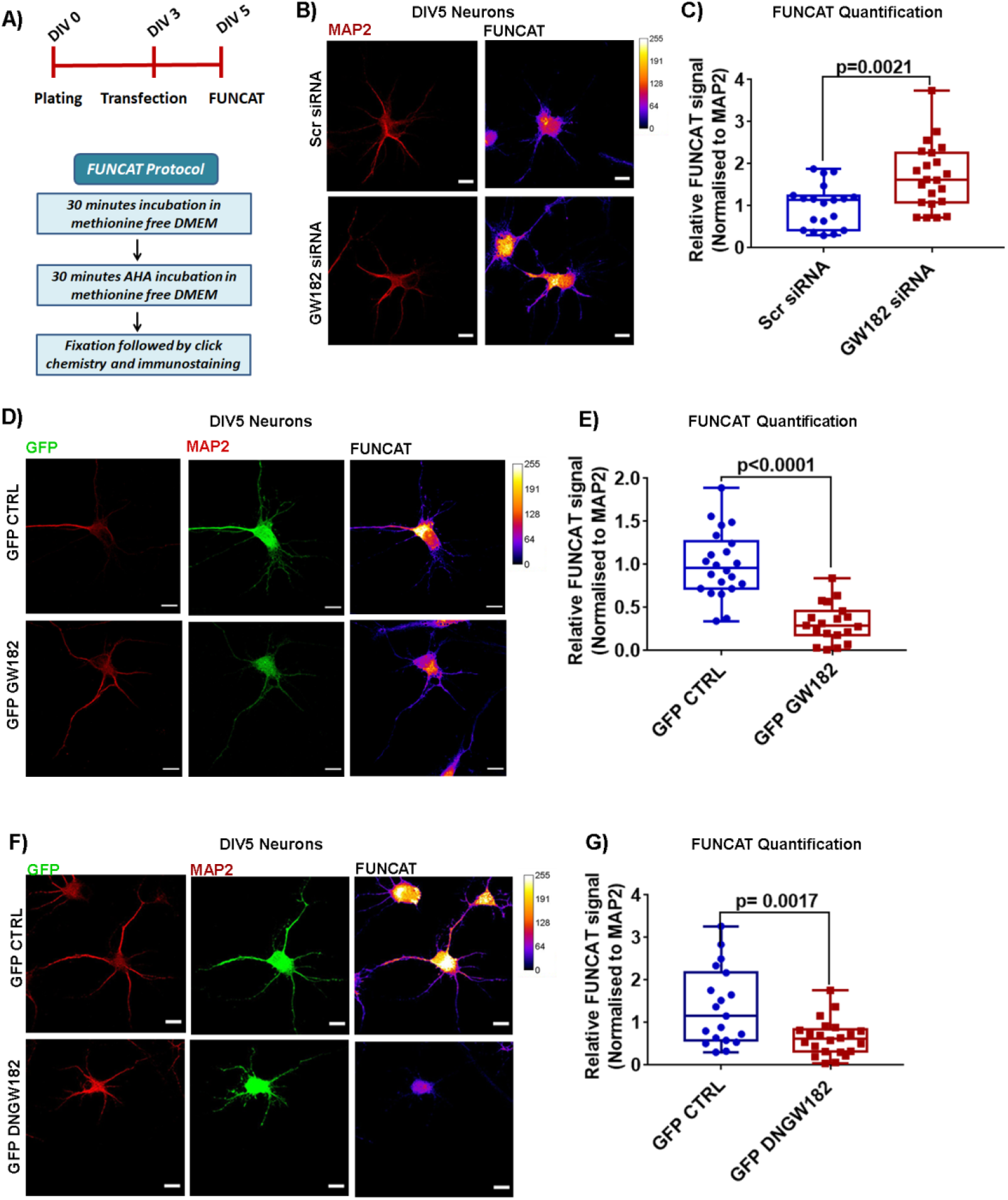
GW182 regulates Global translation in developing neurons. A. Schematic showing the experimental procedure and timeline to visualize and quantify global proteins synthesis in DIV5 cultured hippocampal neurons through the fluorescent non-canonical amino acid tagging metabolic labeling technique (FUNCAT). B. Representative images of DIV5 cultured hippocampal neurons showing FUNCAT signal in Scrambled siRNA or GW182 siRNA transfected neurons, Scale bar represents 50 microns. C. Quantification of normalized FUNCAT signal (Normalized to corresponding MAP2 signal) in Scrambled siRNA or GW182 siRNA transfected neurons. n=19-22 neurons from 3 independent experiments, Unpaired t-test with Welch’s correction. D. Representative images of DIV7 cultured hippocampal neurons showing FUNCAT signal in control GFP and GFP GW182 transfected neurons, Scale bar represents 50 microns. E. Quantification of FUNCAT signal (Normalized to corresponding MAP2 signal) in control GFP and GFP GW182 transfected neurons. n=22-26 neurons from 3 independent experiments, Unpaired t-test with Welch’s correction. F. Representative images of DIV7 cultured Hippocampal neurons showing FUNCAT staining in control GFP or GFP-DNGW182 transfected neurons, Scale bar represents 50 microns. G. Quantification of FUNCAT signal (Normalized to corresponding MAP2 signal) in control GFP or GFP DNGW182 transfected neurons, n=21 neurons from 2 independent experiments, Unpaired t-test with Welch’s correction.

We found that GW182 knockdown resulted in over 70% increase in somatodendritic FUNCAT intensity of neurons at DIV5 **(Figure 6B and 6C)**. However, when we examined the effect of GW182 knockdown in neurons at DIV6, we did not observe any significant difference in the somatodendritic FUNCAT signal **(Figure S6B and S6C)**. Conversely, GW182 overexpressing neurons showed over 70% reduction in somatodendritic FUNCAT intensity at DIV5 **(Figure 6C and 6D)**.

In addition, we also tested the effect of DN GW182 on neuronal translation. In our morphology experiments, we found that DNGW182 influences dendritic morphology in a similar manner to that of GW182 knockdown (figure 3 and figure 4). Surprisingly, we found that in contrast to increased FUNCAT signal in GW182 knockdown cells, overexpression of DNGW182 resulted in reduced somatodendritic FUNCAT signal in cultured neurons at DIV5 and DIV6 **(Figure 6E, 6F, S6C, and S6D)**. This indicated that overexpression of DN GW182 is not the same as knockdown of GW182 with respect to translation response. Overall, we found that GW182 downregulates global translation in the neuronal somatodendritic compartment of developing neurons.

### GW182 is a crucial regulator of dendritic F-actin and LIMK1 levels

In the previous set of experiments, we clearly established the role of GW182 in the regulation of protein synthesis and dendritic arborization. Since cytoskeletal remodeling is a fundamental determinant of dendritic arborization, we hypothesized that GW182 might impact dendritic architecture via regulation of neuronal cytoskeleton (Jan and Jan, 2010). Both actin and microtubule elements play an essential role in determining the dendritic architecture. Here we have focused to study if GW182 regulates dendritic actin dynamics. Actin is known to exist in two states: the monomeric globular-actin (G-actin), which polymerizes to form asymmetric two-stranded helical F-actin. Actin polymerization dynamics (actin treadmilling; F-actin/G-actin ratio) regulate distinct features of dendritic morphology (Georges et al., 2008; Ravindran et al., 2019; Wolterhoff et al., 2020). Here, we determined the influence of GW182 expression on dendritic F-actin levels. We measured the amount of dendritic F-actin using Phalloidin staining. GW182 expression was modified by transfecting the neurons with Scrambled siRNA/GW182 siRNA, or GFP/GFP GW182/GFP DN GW182 at DIV3, followed by fixing the neurons at DIV7 and immunostaining with Phalloidin and MAP2 **(Figure7A)**. Neurons transfected with GW182 siRNA demonstrated significantly increased levels of dendritic F-actin **(Figure 7B and 7C)**. Conversely, neurons with GFP-GW182 overexpression showed reduced F-actin levels in the dendrites **(Figure 7D and 7E)**. Surprisingly, with the overexpression of DNGW182, we did not see any change in the dendritic F-actin levels **(Figure 7F and 7G)**. The opposite effect of knockdown and overexpression of GW182 on dendritic F-actin levels indicated that GW182 has an important role in determining dendritic actin dynamics. Currently, we do not have an explanation regarding the F-actin phenotype of DNGW182 overexpressing cells. However, taken together with FUNCAT results of Dominant negative mutant, this shows that the mutant does not behave like GW182 knockdown in every aspect and should be used with caution.

**Figure 7:**
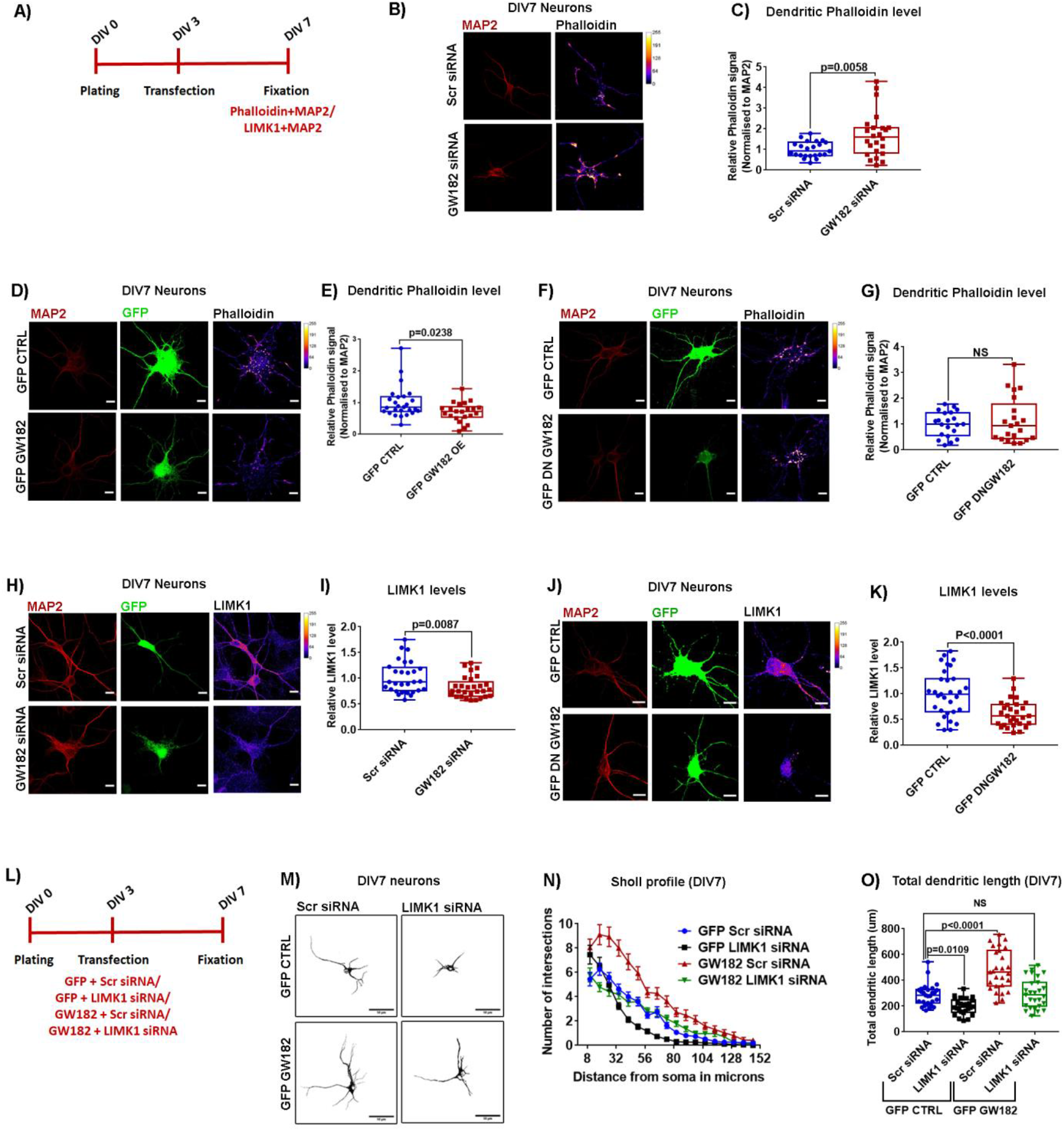
GW182 regulates dendritic arborization through LIMK1 mediated modulation of dendritic F-actin levels. A) Schematic showing the experimental procedure and timeline: Cultured hippocampal neurons were transfected with indicated expression constructs at DIV3 and fixed at DIV7, followed by MAP2 and Phalloidin staining. B) Representative images of DIV7 cultured hippocampal neurons showing Phalloidin staining in Scrambled siRNA or GW182 siRNA transfected neurons. Scale bar represents 10 microns. C) Quantification of Phalloidin intensity (Normalized to corresponding MAP2 signal) in Scrambled siRNA or GW182 siRNA transfected neurons. n=23-24 neurons from 3 independent experiments, Unpaired t-test with Welch’s correction. D) Representative images of DIV7 cultured hippocampal neurons showing Phalloidin staining in either GFP control or GFP GW182 transfected neurons. Scale bar represents 10 microns. E) Quantification of Phalloidin intensity (Normalized to corresponding MAP2 signal) in GFP control or GFP GW182 transfected neurons. n=22-26 neurons from 3 independent experiments, Unpaired t-test with Welch’s correction. F) Representative images of DIV7 cultured hippocampal neurons showing Phalloidin staining in GFP control or GFP DNGW182 transfected neurons. Scale bar represents 10 microns. G) Quantification of Phalloidin intensity (Normalized to corresponding MAP2 signal) in GFP control or GFP DNGW182 transfected neurons, n=21 neurons from 3 independent experiments, Unpaired t-test with Welch’s correction. H) Representative images of DIV7 cultured hippocampal neurons showing LIMK1 staining in scrambled siRNA or GW182 siRNA transfected neurons. Scale bar represents 10 microns. I) Quantification of normalized LIMK1 signal in Scrambled siRNA or GW182 siRNA transfected neurons. n=36-39 neurons from 4 independent experiments, Unpaired t-test with Welch’s correction J) Representative images of DIV7 cultured hippocampal neurons showing LIMK1 staining in GFP control or GFP DNGW182 transfected neurons. Scale bar represents 10 microns. K) Quantification of normalized LIMK1 signal in GFP control or GFP DNGW182 transfected neurons. n=29-31 neurons from 4 independent experiments, Unpaired t-test with Welch’s correction. L) Schematic showing the experimental procedure and timeline: Cultured hippocampal neurons were transfected with indicated expression constructs at DIV3 and fixed at DIV7, followed by MAP2 immunostaining. M) Representative micrographs of DIV7 cultured hippocampal neurons transfected with control GFP or GFP GW182 along with scrambled siRNA or LIMK1 siRNA. The images are derived from the threshold MAP2 intensity of transfected neurons. Scale bar represents 50 microns. N) Intersection profile / Sholl curve of DIV7 cultured hippocampal neurons transfected with GFP GW182 or control GFP along with scrambled siRNA or LIMK1 siRNA. Data: mean +/- SEM, n=32 neurons from 4 independent experiments, GFP + LIMK1 siRNA transfected neurons had significantly less number of dendritic intersection than GFP+ Scrambled siRNA transfected neurons at 8, 32-64 microns from the soma, GFP GW182+ Scrambled siRNA overexpressing neurons had significantly more dendrites from GFP+ Scrambled siRNA overexpressing neurons at 16-88 microns from the soma, and GFP GW182+ LIMK1 siRNA overexpressing neurons had significantly more dendrites from GFP+ Scrambled siRNA overexpressing neurons only at 16 microns from the soma, two way ANOVA followed by Bonferroni’s multiple comparisons test. O) Quantification of the total dendritic length of cultured hippocampal neurons transfected with either control GFP or GFP GW182 along with scrambled siRNA or LIMK1 siRNA. n=26-27 neurons from 3 independent experiments, One way ANOVA followed by Bonferroni’s multiple comparisons test.

We next sought to identify the mechanism behind GW182 mediated regulation of dendritic Factin levels. Multiple Actin binding proteins cooperate in controlling the structure and stability of the neuronal actin network (Jan and Jan, 2010). Among these, LIMK1 acts as a key regulatory protein, known to control dendritic morphology via regulation of actin polymerization (Ravindran et al., 2019; Saito et al., 2013). Furthermore, LIMK1 is translationally regulated in neurons downstream of external cues and is also regulated by miRNA machinery (Ravindran et al., 2019; Schratt et al., 2006). We investigated whether an alteration in GW182 levels influences LIMK1 expression in neurons. To address this, we undertook a loss of function approach. We performed GW182 knockdown in culture hippocampal neurons at DIV3 by transfecting the neurons with scrambled siRNA or GW182 siRNA along with GFP. The transfected neurons were fixed at DIV7 and examined for the levels for LIMK1 using quantitative immunofluorescence. LIMK1 levels were significantly reduced in neurons with GW182 knockdown as compared to the neurons transfected with Scrambled siRNA **(Figure 7H and 7I)**.

We further validated this result by employing DN GW182 mutant to perturb the GW182 function. We quantified LIMK1 levels in DIV7 neurons transfected with either GFP or GFP DNGW182 construct on DIV3 **(Figure 7A).** Similar to GW182 knockdown, we found that perturbation of GW182 function using dominant DNGW182 also leads to a significant reduction in somatodendritic LIMK1 levels **(Figure 7J and 7K)**. Conversely, we show that overexpression of GW182 leads to increased LIMK1 expression in HEK293T cells **(Figure S7A and S7B)**.

Hence, we have demonstrated that modulation of GW182 expression results in altered LIMK1 and F-actin levels. We speculated that GW182 regulates dendritic arborization through modulation of LIMK1 levels. In that case, knocking down LIMK1 should limit the dendritic overgrowth phenotype caused by GW182 overexpression.LIMK1 siRNA was validated using immunostaining analysis in hippocampal neurons **(Figure S7C and S7D).** After initial validation of LIMK1 siRNA, to test our hypothesis, we co-transfected DIV3 hippocampal neurons with Scrambled or LIMK1 siRNA in the background of GFP/GFP GW182 overexpression **(Figure 7L)**. These neurons were fixed at DIV7 and immunostained with MAP2 antibody. In agreement with our previous result, neurons with GFP GW182 and scrambled siRNA expression showed increased dendritic arborization as compared to neurons with GFP control and Scrambled siRNA expression **(Figure 7M and 7N**). In GFP transfected neurons, knocking down LIMK1 resulted in a significant reduction in dendritic branching and total dendritic length **(Figure 7M, 7N, and 7O)**. Importantly, LIMK1 knockdown partially rescued the dendritic overgrowth phenotype of GW182 overexpression **(Figure 7M and 7N**). We also observed that LIMK1 siRNA transfection prevented GW182 induced increase in total dendrite length and total dendritic intersections **(Figure 7O and S7E**). Taken together, these results suggest that GW182 regulates dendritic morphology via LIMK1 mediated modulation of dendritic actin dynamics.

## Discussion

Till now, the research on GW182 has been primarily focused on understanding its role as a miRISC component. In our study, we found a strictly regulated expression pattern of GW182 during neuronal development, with specific periods of peak expression and decline. The period of peak GW182 expression coincided with the period of rapid dendritogenesis, followed by reduced GW182 expression in the adult brain. In the hippocampus, GW182 expression peaked at P7, followed by a substantial reduction from P15 onwards. In the cerebellum, the peak of GW182 levels was observed at P15, with a significant reduction at P30, followed by a further reduced level at P60. The temporal delay in peak and drop of GW182 expression in the cerebellum is likely due to the delayed development of cerebellar circuitry. Immunohistochemical analysis of GW182 in hippocampal and cerebellar sections revealed the presence of GW182 in a wide variety of neuronal cells including glutamatergic (Pyramidal and Granule cells) as well as GABAergic neurons (Purkinje neurons). During its period of peak expression, GW182 staining was observed in the dendritic region of both hippocampal and cerebellar neuronal populations. Furthermore, later during development, the reduction in GW182 staining was predominantly observed in the cyto-dendritic compartment. The presence of GW182 in the dendritic compartment during the period of peak GW182 expression indicates important dendritic functions for GW182. Interestingly, our imaging analysis also revealed substantial staining of GW182 in the neuronal nucleus. As previous studies have indicated an important role of nuclear GW182 in RNA-mediated transcriptional activation (Hicks et al., 2017), it is important to determine the nuclear function of GW182 in neurons.

Whether the expression of GW182 is regulated during neuronal development by transcription or translation or turnover needs to be explored further. We observed that the amount of GW182 mRNA remains unchanged during neuronal development, suggesting post-transcriptional regulation of GW182 expression during neuronal development. Studies have indicated that GW182 levels can be regulated post-transcriptionally as well as through ubiquitin-mediated proteasomal degradation (Li et al., 2014; Olejniczak et al., 2016). Our expression profile studies indicated that the developmental reduction in GW182 expression correlates with the emergence of synaptic circuitry and neuronal activity. This persuaded us to explore the potential contribution of neuronal activity in controlling GW182 levels. To elucidate this, we modulated the neuronal activity in hippocampal cultures using tetradotoxin or picrotoxin. Tetradotoxin is a sodium channel blocker which leads to a reduction in neuronal activity. Application of tetradotoxin increased GW182 expression. Alternatively, when we increased the neuronal activity in our culture system by using GABA antagonist picrotoxin, we observed a reduction in GW182 levels. Taken together, these results indicate the potential role of neuronal activity in the regulation of GW182 expression. Many previous studies have demonstrated an important role of PI3K-Akt-mTOR and JAK-STAT pathways in regulating the translation of GW182 (La Rocca et al., 2015; Olejniczak et al., 2016). It is important to determine if these pathways are also involved in neuronal activity mediated regulation of GW182.

Abundance of GW182 in dendrites during the period of rapid dendritogenesis indicates its role in dendrite morphogenesis. To understand the role of GW182 in dendritic morphogenesis, we perturbed GW182 function during the dendritic growth phase of hippocampal neurons using two independent loss of function approaches; siRNA knockdown and expression of Dominant Negative (DN) GW182. Both these approaches resulted in the reduction of dendritic arborization. Consistent with this, overexpression of GW182 resulted in increased dendritic arborization. These results establish GW182 as a positive modulator of dendritic growth. Importantly, we observed that GW182 regulates dendritic arborization only during a specific temporal window, as loss of function of GW182 did not affect dendritic arborization in older neurons (DIV7-11). We hypothesize that this is mainly due to low basal levels GW182 at this stage. We also observed that perturbation of GW182 function during DIV 3-7 did not affect BDNF induced dendritic growth. This is in contrast to a previous study which suggested that loss of GW182 prevents BDNF induced dendritic growth in mature neurons (DIV14) (Huang et al., 2012). We speculate that these contrary findings are primarily due to the differences in the age of neurons and the basal expression of GW182 during the temporal stages examined. Altogether, these results establish GW182 as an important modulator of dendritic growth.

GW182 is likely to regulate dendritic arborization by modulating cytoskeletal elements such as actin and microtubule. The actin cytoskeletal organization is a crucial determinant of dendritic architecture. Consequently, multiple factors determining dendritic arborization converge on controlling the activity and polymerization actin cytoskeleton (Konietzny et al., 2017). Here, we show that GW182 regulates dendritic F-actin levels. Our experiments revealed reduced dendritic F-actin levels in neurons with GW182 overexpression and increased F-actin staining in GW182 knockdown neurons. Although the exact relation between dendritic F-actin levels and dendritic arborization remain contradictory, substantial evidence has suggested that destabilization of actin filaments in dendrites is required for microtubule invasion of the dendritic filopodia, which in turn leads to branch stabilization and increase in dendritic length (Poulain and Sobel, 2010; Ravindran et al., 2019). Hence, reduced F-actin levels observed in GW182 overexpressing neurons correlates well with the increased dendritic arborization of these neurons. These experiments establish the actin cytoskeleton as an important player in GW182 mediated dendritic growth. However, at this stage, we cannot rule out the potential contribution of the microtubule cytoskeleton in this regulation.

Previous reports have demonstrated the important role of translation regulation in dendritic morphogenesis (Chihara et al., 2007; Kumar et al., 2005; Lein and Higgins, 1991; Perycz et al., 2011). In most studies, an increase in protein synthesis leads to increased dendritic arborization and vice-versa (Keil et al., 2018; Kumar et al., 2005; Skalecka et al., 2016; Xu et al., 2019). However, in our study, we found a new mechanism where despite causing disinhibition of global protein synthesis, GW182 knockdown resulted in reduced dendritic arborization. We speculated that in the background of global translation up-regulation, knockdown of GW182 might lead to the translation down-regulation of selective mRNA candidates required for dendritic growth. We have identified actin regulator LIMK1 as one of such candidates whose expression is selectively downregulated despite global translation up-regulation observed in GW182 knockdown. In accordance with a previous study from our lab, we observed that the dynamic shift in actin polymerization was well correlated with somatodendritic LIMK1 levels upon modulation of GW182 expression (Ravindran et al., 2019). Furthermore, we showed that LIMK1 knockdown could rescue the dendritic overgrowth phenotype of GW182 overexpression. In summary, we showed that GW182 regulates dendritic arborization via LIMK1 induced changes in the dendritic actin cytoskeleton **(Figure 8)**. An important direction for future studies is to determine the molecular mechanism of GW182 regulation of LIMK1 expression.

**Figure 8:**
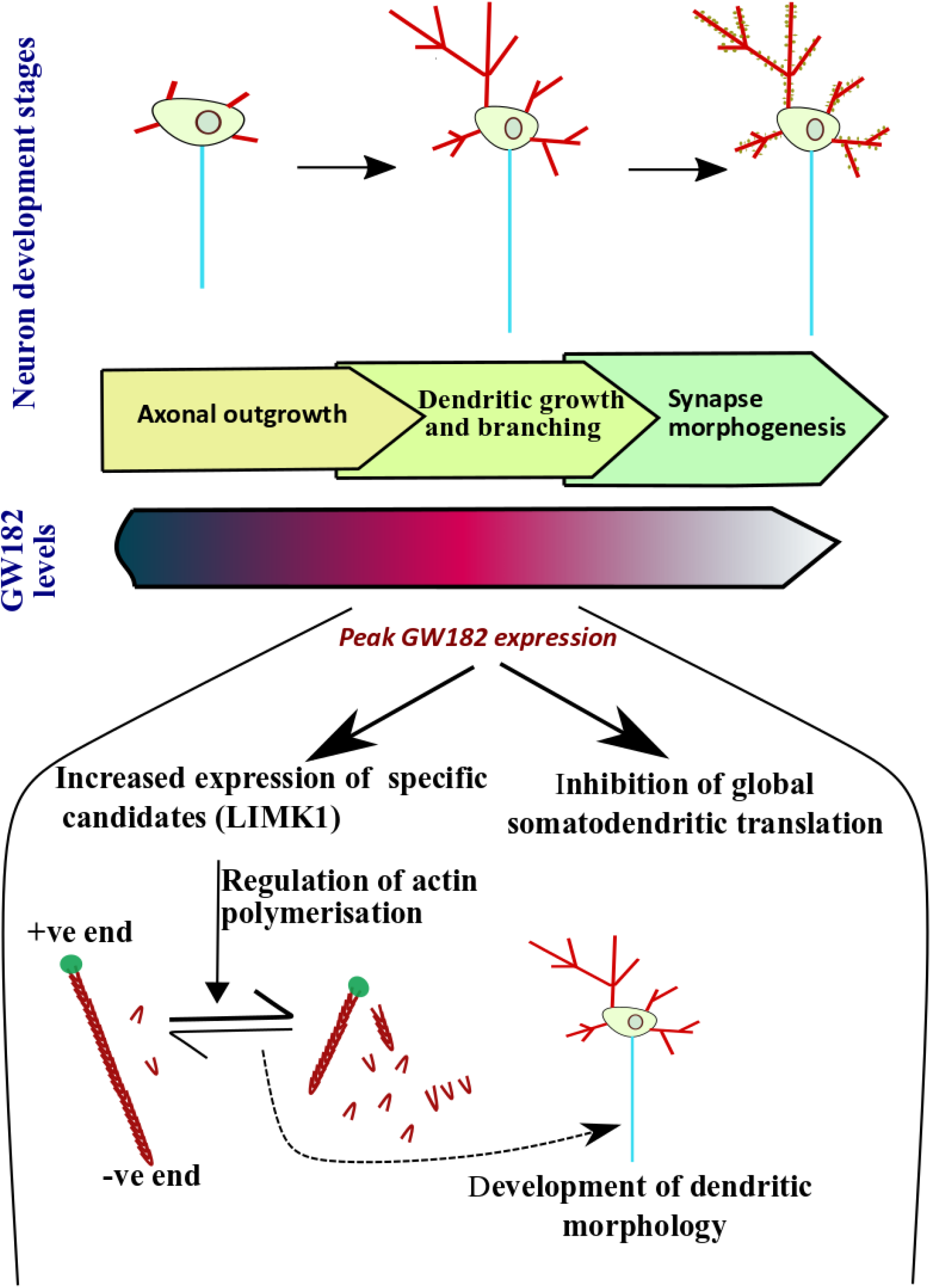
Model describing GW182 mediated regulation of dendritic arborization. ***A)*** Peak expression of GW182 during the period of rapid dendritogenesis regulates dendritic arborization via LIMK1 mediated regulation of actin polymerization.

In this study and many previous studies, people have used DNGW182 as a substituent for GW182 knockdown (Huang et al., 2012; Jakymiw et al., 2005). In our dendritic morphology assay, DNGW182 behaved similarly to GW182 knockdown leading to a reduction in dendritic arborization. However, in FUNCAT assay, in contrast to the translation up-regulation phenotype of GW182 knockdown, overexpression of DNGW182 resulted in translation inhibition. We speculate that the DN GW182 causes translation inhibition by sequestering the AGO2-miRNA-mRNA complex irreversibly, thereby preventing mRNA translation. Furthermore, in our experiments involving the measurement of dendritic F-actin levels, the DNGW182 did not phenocopy GW182 knockdown. Taken together, these results suggest DNGW182 mutant does not behave like GW182 knockdown in every aspect and should be used with caution.

In conclusion, our work established the crucial role of GW182 in regulating dendritic morphogenesis during a restricted developmental window. As GW182 has recently been implicated in Neurodevelopmental disorder as well as Alzheimer’s disease (Badhwar et al., 2017; Chen et al., 2019; Eising et al., 2019; Guerrini and Mei, 2018), our work provides a framework for future studies delineating the role of GW182 in different aspect of neuronal development and function. Important questions to be addressed in the future includes a better understanding of the mechanism behind GW182 mediated dendritic development, electrophysiological, and behavioral characterization of circuits/ animals with alteration in GW182 expression. Furthermore, we speculate that the low levels of GW182 in mature neurons are important in providing reversibility to miRISC and in determining translation response to various neurotransmitter stimuli.

## Materials and methods

### Ethics Statement

All animal work was carried out in accordance with the procedures approved by the Institutional Animal Ethics Committee (IAEC) and the Institutional Biosafety Committee (IBSC), InStem, Bangalore, India. All rodent work was done with Sprague Dawley (SD) rats. Rats were kept in 20–22 °C temperature, 50–60 relative humidity, 0.3 μm HEPA-filtered air supplied at 15–20 ACPH, and 14-h/10-h light/dark cycle maintained. Rats were freely fed with Food and water.

### Cell lines and primary neuronal culture

Primary neuronal cultures were prepared from the hippocampus of Sprague-Dawley rats at embryonic day 18 (E18) as per the previously established procedure (Kaech and Banker, 2006; Ravindran et al., 2019). The dissociated cells were plated at a density of 20000-30000 cells/cm^2^ on poly-L-lysine (P2636, Sigma) (0.2mg/ml in borate buffer, pH 8.5) coated dishes/coverslips. The neurons were initially plated in Minimum Essential Media (MEM, 10095080, Thermo Fisher Scientific) containing 10% FBS (F2442, Sigma) to aid their attachment. After 3hours, the media was changed to neurobasal (21103049, Thermo Fisher Scientific) supplemented with B27 (17504044, Thermo Fisher Scientific) and Glutamax (35050-061, Life Technologies). Neurons were cultured for the required time at 37 °C in a 5% CO_2_ incubator. Transfections were done using Lipofectamine 2000 (11668019, Invitrogen) following modified manufacture’s protocol. Briefly, 2ug of DNAor100pmol of siRNA was used along with 4ul of L2000 for per well of a 6 well dish. The cells were incubated in the transfection mixture for 2hr followed by media change.

HEK293T and Neuro2A cells were maintained in DMEM (11995, Gibco) with 10% FBS. Cells were cultured for the required time at 37°C in a 5% CO_2_ incubator.

### Lysate preparation and immunoblotting

Rats of different postnatal ages were sacrificed and required tissue (hippocampus, cerebellum, and liver) was dissected out in cold Phosphate Buffered Saline (PBS) of pH 7.4. The tissue was homogenized in lysis buffer containing 50 mM Tris pH 7.4, 150mM NaCl, 5mM MgCl2, 0.3% Triton X-100, supplemented with EDTA-free protease inhibitor complex (Sigma, cat. no. S8830) and phosphatase inhibitor cocktail (04906837001, Roche). The homogenates were centrifuged at 16000 RCF for 30 min at 4°C. Obtained lysates were mixed with 6X laemmli buffer and were heated at 95 °C. The samples were aliquoted and stored at −20°C until further use. Prepared lysates were loaded onto SDS PAGE gel and ran for 180 minutes. For experiments involving immunoblotting of GW182, a 6% resolving gel was used. In the rest of the experiments, 8% or 10% resolving gel was used depending upon the molecular weight of proteins to be resolved. Proteins were then transferred to a PVDF membrane (Overnight transfer at 20 V/ 2hrs transfer at 380mA). Post transfer, the blots were blocked with 5% blotto or 5% BSA (for phospho antibodies) for 1hr at room temperature. Afterward, the blots were incubated with the required primary antibody for 3hrs at room temperature. The blots were washed with 1% TBST (TBS+ Tween 20; GRM156, HIMEDIA) thrice for 10 min each, and incubated with appropriate secondary antibodies for 1hr at room temperature. After subsequent washes with TBST, the blots were developed by a chemiluminescent method using ECL western clarity solution (1705060, Biorad). Images were taken in Image Quant (LAS 4000 / Amersham imager 600). The bands were quantified using Image J software.

### RNA extraction and Quantitative PCR

Total RNA was extracted from the homogenized lysates by Trizol (15596026, Thermo Fisher Scientific) method and isolated RNA was converted to cDNA using superscript III (18080, Invitrogen). Quantitative PCR (qPCR) was performed for required primers using Biorad thermal cycler CFX384.

List of primers

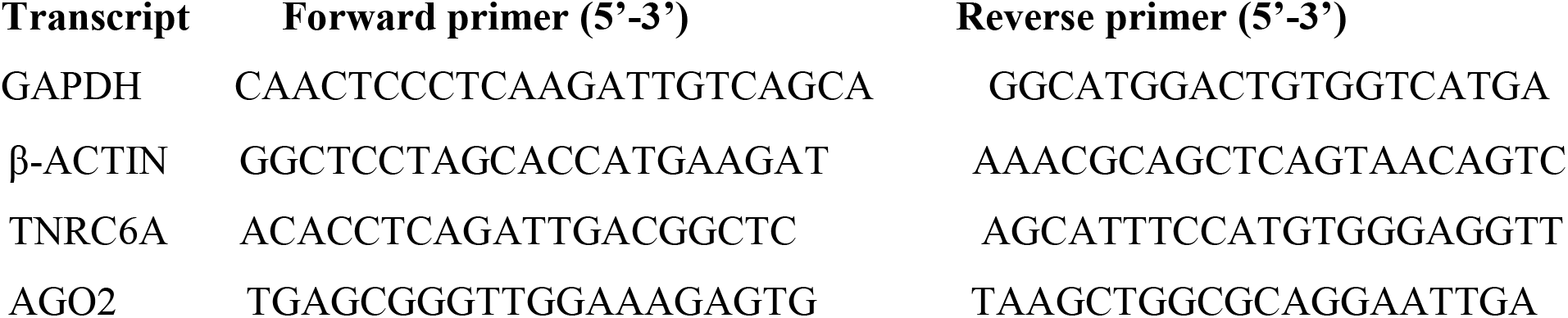

### Immunostaining

Rat primary hippocampal neurons were fixed with 4% PFA for 20 minutes at room temperature and processed for imaging as described before (Muddashetty et al., 2011). In brief, cells were permeabilized using TBS_50_T (0.3%) [50 mM Tris-Cl (pH 7.4) + 150 mM NaCl + 0.3% Triton X-100]. Afterwards, the cells were blocked with blocking buffer [TBS_50_T (0.1%) + 2% BSA + 2% FBS] for 1 hour at room temperature. Neurons were incubated with required antibodies for two hours at room temperature, followed by washes with TBS_50_ T (0.1%). After washes, neurons were incubated with suitable secondary antibodies for 1 hour at room temperature. After washes, the coverslips were mounted for imaging using Mowiol^®^ 4-88 mounting media. Images were acquired on FV3000 confocal microscope (Olympus). For Neuronal morphology experiments involving Sholl analysis and length measurements, 40X NA 1.25, silicon oil immersion objective was used, along with 1μm step size in the z-direction. For other experiments involving the quantification of fluorescent intensities, images were taken at 60X, NA 1.4, oil immersion objective with 0.5μm step size in Z-axis. Imaging conditions were kept constant across different data sets in an experiment. In all the experiments, the pinhole was kept at 1 Airy unit.

### F-actin measurement

For visualization of F-actin, Alexa Fluor 488 phalloidin (A12379, Invitrogen) was added to the secondary antibody solution (1:50 dilution) during immunostaining and incubated for 1 h. Only the dendritic F-actin levels were measured for intensity calculation. The F-actin levels were quantified as an intensity ratio of phalloidin to Map2 fluorescence. Image analysis was performed using FIJI software and the maximum intensity projection of the slices was used for quantification of the mean fluorescent intensities. The mean fluorescent intensity of the dendritic Phalloidin signal was normalized to the corresponding MAP2 intensity.

### FUNCAT (Fluorescent non-canonical amino acid tagging)

For metabolic labeling, cultured hippocampal neurons were incubated in Methionine-free DMEM for 30 minutes. Afterward, the neurons were treated with L-azidohomoalanine **(AHA, 1μM)** (1066100, Click Chemistry tools) for 30 minutes in Met-free DMEM (21013024, Thermo Fisher Scientific). The coverslips were then washed once with 1X PBS and fixed with 4% PFA for 20 minutes at room temperature. After fixation, the neurons were permeabilized for 10 minutes with 0.3% Triton X-100 solution prepared in TBS_50_ (50mM Tris, 150mM NaCl, pH 7.6). The permeabilized neurons were blocked in blocking buffer [TBS_50_T (0.1%) + 2% BSA + 2% FBS] for 1 hour. Afterward, the neurons were subjected to the FUNCAT reaction for 2 hours where the newly-synthesized AHA incorporated proteins were tagged with an alkyne-fluorophore Alexa-Fluor 555 through click reaction (C10269, CLICK-iT cell reaction buffer kit, Click Chemistry Tools). After 3 washes with TBS_50_T (0.1%), the neurons were immunostained with MAP2 antibody, followed by mounting with mowiol medium. The cells were imaged on Olympus FV300 confocal laser scanning inverted microscope with 60X objective. The pinhole was kept at 1 Airy Unit and the objective was moved in Z-direction with a step size of 0.5μM to collect light from the planes above and below the focal plane. Image analysis was performed using FIJI software and the maximum intensity projection of the slices was used for quantification of the mean fluorescent intensities. The mean fluorescent intensity of the FUNCAT signal was normalized to the MAP2 intensity of the corresponding neuron.

### Perfusion and sectioning

SD rats of required postnatal ages were transcardially perfused with 4% paraformaldehyde (PFA). Afterward, the brain was carefully dissected out of the skull and kept in 4% PFA overnight for post-fixation. Fixed tissue was washed thrice with 0.1M phosphate buffer (PB) and stored at 4 °C until sectioning. 50 micron thick sections were obtained from the brain required region using Leica VT 1200S vibrating blade microtome. The sections were used for further immunohistochemistry as described above.

### Image analysis

All the image analysis was done using Fiji (ImageJ-based image processing package) software. For quantification of fluorescence intensity, the confocal stacks were collapsed using the maximum intensity projection method, and the mean intensity was quantified from the cells. the mean fluorescent intensity of the signal was normalized to the corresponding MAP2 mean fluorescent intensity. For sholl analysis quantification, the confocal stacks were first collapsed using the maximum intensity projection method. Thereafter, the MAP2 channel of the images was thresholded manually and this was used for carrying out sholl analysis using the sholl plugin in ImageJ. In all analyses, the starting radius was kept as 10 microns with a step size of 8 microns. For measurement of dendritic length, thresholded MAP2 images were traced using the NeuronJ plugin in ImageJ software as described (Meijering et al., 2004).

### Statistical analysis

Statistical analysis was performed using Graph pad prism software. Data distribution was tested for normality using Kolmogorov Shapiro Smirnov goodness-of-fit test. Depending on the distribution, either parametric or non-parametric tests were used to quantify statistical significance. For groups with less than 5 data points, we assumed the data to be normally distributed. For comparing two groups, Student’s t-test (two-tailed, unpaired) was used for normally distributed data with equal variance, Student’s t-test with Welch’s correction was used for normally distributed data with unequal variances, and Mann-Whitney test was used to compare data with non-normal distribution. Multiple group comparisons were made using oneway ANOVA followed by Bonferroni’s multiple comparisons test. Sholl profile between two different groups was determined using Two-way ANOVA followed by Bonferroni’s multiple comparisons test. *P* values of less than 0.05 were considered statistically significant. Data are presented as mean ± Standard error of Mean (SEM).

### Antibodies, plasmids, and other reagents

#### Antibodies

GW182 (G5922, Sigma), AGO2 (H00027161-MO1, Abnova), FMRP (F4055, Sigma), Map2 (M9942, Sigma), MOV10 (ab80613, Abcam), Calbindin (214005, Synaptic Systems), Tuj1 (T8578, Sigma), XRN1 (SAB4200028, Sigma), LIMK1 (ab81046, Abcam), ERK (9102, Cell Signaling Technologies), p-ERK (9101, Cell Signaling Technologies), GFP(ab6556,Abcam) Secondary Rabbit HRP (A0545, Sigma), Secondary Mouse HRP (31430, Thermo Fisher Scientific),α tubulin (T9026, Sigma), Alexa Fluor 488 (A-11059, Thermo Fisher Scientific), Alexa Fluor 555 (A-21428, Thermo Fisher Scientific), Alexa flour 647 (A-21235, Thermo Fisher Scientific).

#### Plasmids and siRNA

Limk1 siRNA (s134717, Thermo Fisher Scientific), GW182 siRNA (s107649, Thermo Fischer Scientific). pmyc-GFP-TNRC6A was a gift from Kumiko Ui-Tei (Addgene plasmid # 41999; http://n2t.net/addgene:41999; RRID: Addgene_41999). GFP-GW182delta1 was a gift from Edward Chan (Addgene plasmid # 11592; http://n2t.net/addgene:11592; RRID: Addgene_11592).

#### Chemicals

Tetradotoxin and picrotoxin were obtained from Tocris Biosciences.

## Acknowledgments

We are thankful to the Central Imaging and Flow-cytometry Facility (CIFF), and the animal house facility of NCBS-instem for providing technical support. We extend our gratitude to all the members of Ravi’s lab for invaluable discussions and suggestions.

## Author contributions

BN designed and performed the experiments, analyzed data, and co-wrote the manuscript, RM conceptualized the project, designed experiments, provided resources, and co-wrote the manuscript.

## Supplementary figure legends

**Figure 1 Supplementary.**
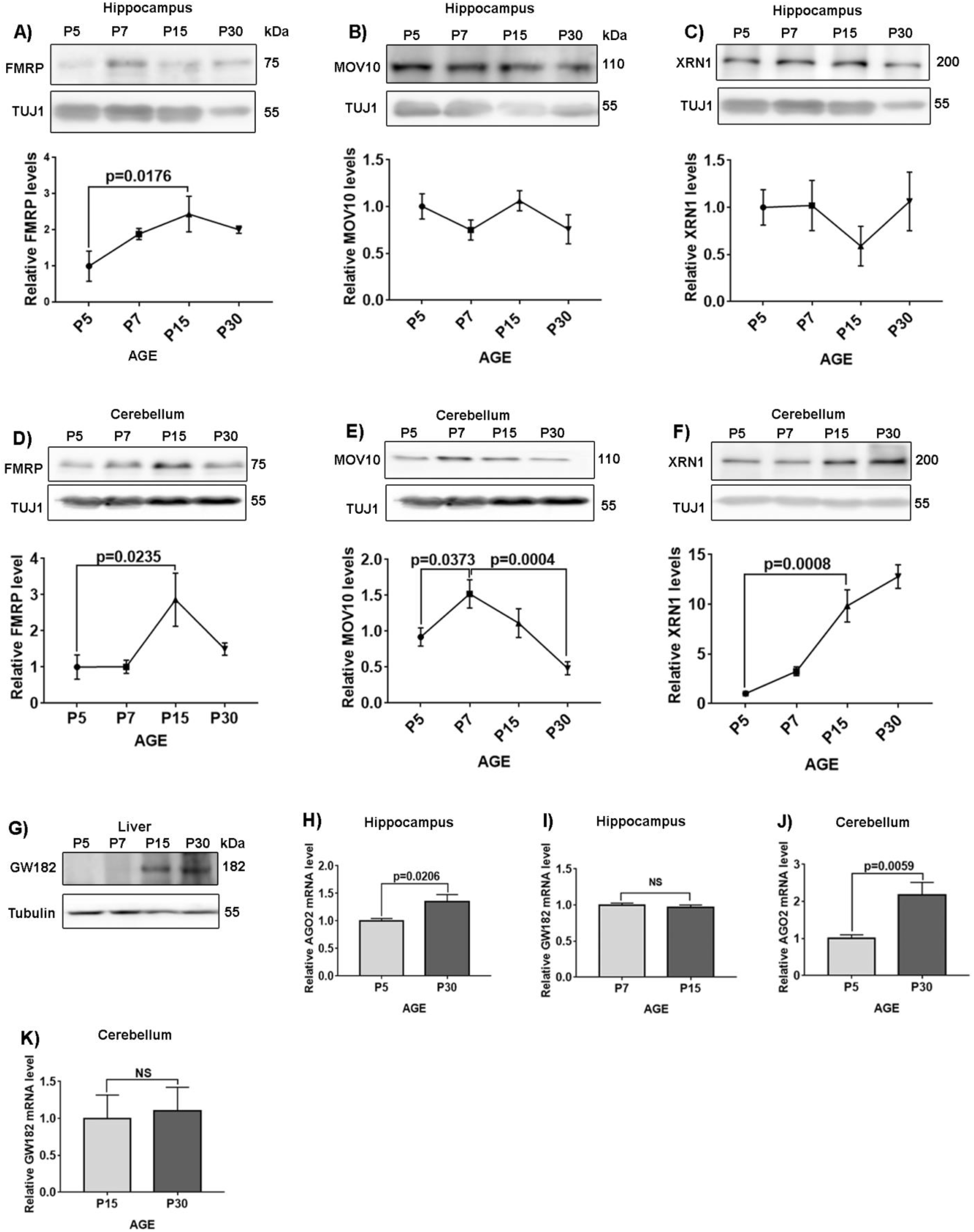
A) Representative immunoblots (top) and line graph (bottom) depicting FMRP expression profile during hippocampal development. Data represent relative FMRP levels normalized to TUJ1, Data: mean +/- SEM, n=3-5 animals per group. One Way ANOVA followed by Bonferroni’s multiple comparisons test. B) Representative immunoblots (top) and line graph (bottom) depicting MOV10 expression profile during hippocampal development. Data represent relative MOV10 levels normalized to TUJ1, Data: mean +/- SEM, n=3-5 animals per group. One Way ANOVA followed by Bonferroni’s multiple comparisons test. C) Representative immunoblots (top) and line graph (bottom) depicting XRN1 expression profile during hippocampal development. Data represent relative XRN1 levels normalized to TUJ1, Data: mean +/- SEM, n=3-5 animals per group. One Way ANOVA followed by Bonferroni’s multiple comparisons test. D) Representative immunoblots (top) and line graph (bottom) depicting FMRP expression profile during Cerebellar development. Data represent relative FMRP levels normalized to TUJ1, Data: mean +/- SEM, n=3-5 animals per group. One Way ANOVA followed by Bonferroni’s multiple comparisons test. E) Representative immunoblots (top) and line graph (bottom) depicting MOV10 expression profile during Cerebellum development. Data represent relative MOV10 levels normalized to TUJ1, Data: mean +/- SEM, n=3-5 animals per group. One Way ANOVA followed by Bonferroni’s multiple comparisons test. F) Representative immunoblots (top) and line graph (bottom) depicting XRN1 expression profile during cerebellum development. Data represent relative XRN1 levels normalized to TUJ1, Data: mean +/- SEM, n=3-5 animals per group. One Way ANOVA followed by Bonferroni’s multiple comparisons test. G) Representative immunoblots depicting GW182 expression profile during liver development. H) Quantitative PCR analysis of AGO2 mRNA expression in P5 and P30 hippocampus. The levels of AGO2 mRNA were normalized using the expression of β-actin mRNA, Data: mean +/- SEM, n=5 independent experiments, Unpaired t-test. I) Quantitative PCR analysis of GW182 mRNA expression in P7 and P15 hippocampus. The levels of GW182 mRNA were normalized using the expression of β-actin mRNA, Data: mean +/- SEM, n=5 independent experiments, Unpaired t-test. J) Quantitative PCR analysis of AGO2 mRNA expression in P5 versus P30 Cerebellum. The levels of AGO2 mRNA were normalized using the expression of β-actin mRNA, Data: mean +/- SEM, n=5 independent experiments, Unpaired t-test. K) Quantitative PCR analysis of GW182 mRNA expression in P15 versus P30 Cerebellum. The levels of GW182 mRNA were normalized using the expression of β-actin mRNA, Data: mean +/- SEM, n=5 independent experiments, Unpaired t-test.

**Figure 2 Supplementary.**
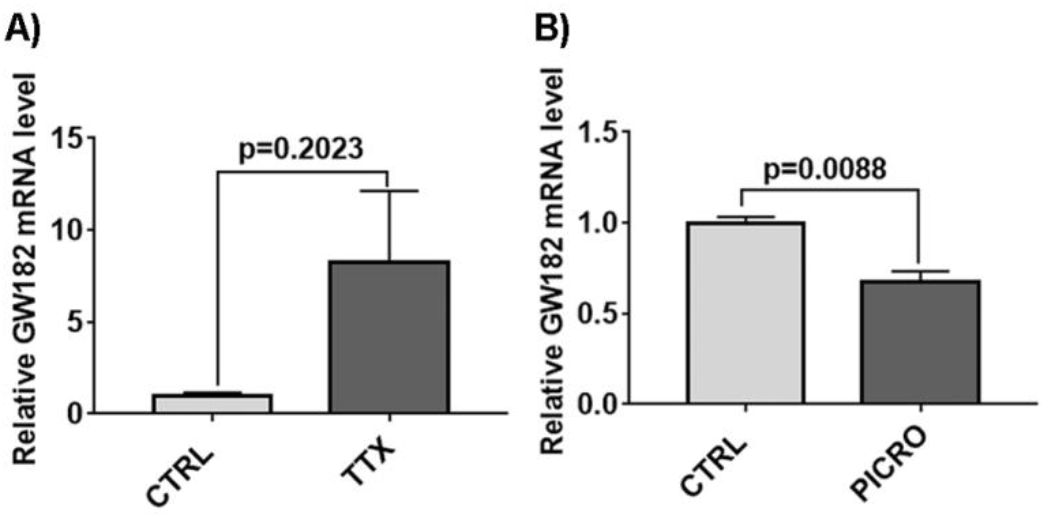
*A)* Quantitative PCR analysis of GW182 mRNA expression from DIV7 hippocampal neurons treated with mock or tetradotoxin for 48hrs. The levels of GW182 mRNA were normalized using the expression of GAPDH mRNA, Data: mean +/- SEM, n=5 independent experiments, Unpaired t-test. *B)* Quantitative PCR analysis of GW182 mRNA expression from DIV7 hippocampal neurons treated with mock or picrotoxin for 48hr. The levels of GW182 mRNA were normalized using the expression of GAPDH mRNA, Data: mean +/- SEM, n=5 independent experiments, Unpaired t-test.

**Figure 3 Supplementary.**
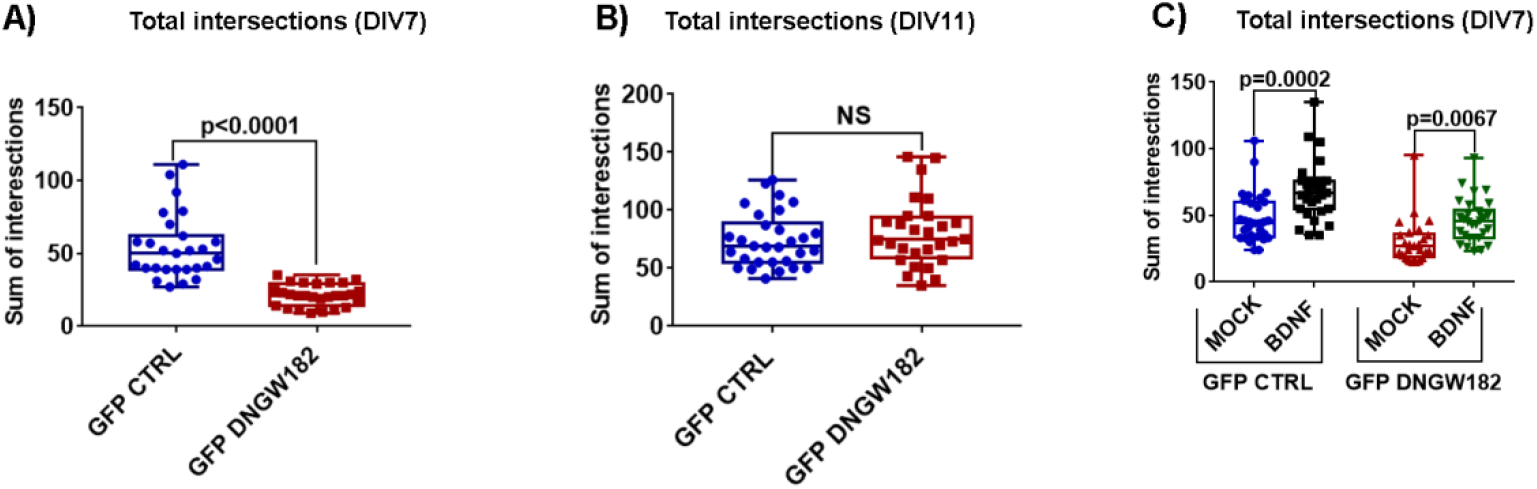
A) Quantification of total intersections of DIV7 hippocampal neurons transfected with either GFP or GFP DNGW182 at DIV 3, n=22-23 neurons from 4 independent cultures, Unpaired t-test with Welch’s correction. B) Quantification of total intersections of MOCK/BDNF treated (48 hrs) DIV7 hippocampal neurons transfected with either GFP or GFP DNGW182, n=22-23 neurons from 4 independent cultures, One Way Anova followed by Bonferroni’s multiple comparison test. C) Quantification of total intersections of DIV11 hippocampal neurons transfected with either GFP or GFP DNGW182 at DIV7, n=22-23 neurons from 4 independent cultures, Unpaired t-test with Welch’s correction

**Figure 4 Supplementary.**
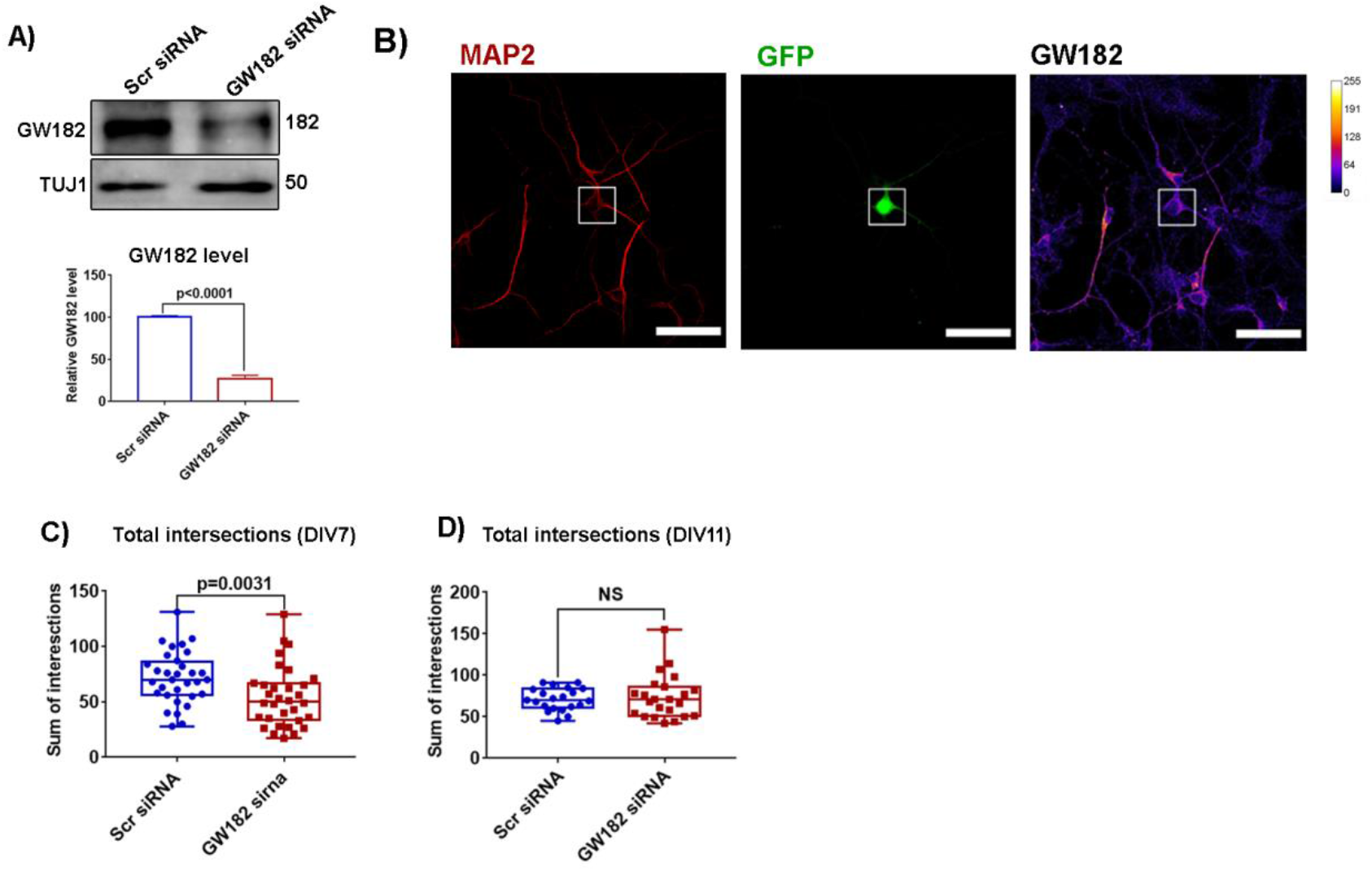
A) Immunoblot validation of GW182 knockdown in Neuro2A cells: Representative immunoblots (top) and quantification (bottom) of GW182 levels from N2A cells treated with either scrambled or GW182 siRNA. Data: mean +/- SEM, n=3 independent experiments, Unpaired t-test. B) Immunostaining validation of GW182 knockdown in hippocampal neuronal culture: Representative image showing GW182 staining in DIV5 hippocampal cultured neurons transfected with GW182 siRNA on DIV3. GFP cotransfection was performed to identify siRNA transfected neurons. ***C)*** Quantification of total intersections of DIV7 hippocampal neurons transfected with either scrambled siRNA or GW182 siRNA on DIV 3, n=22-23 neurons from 4 independent cultures, Unpaired t-test with Welch’s correction. ***D)*** Quantification of total intersections of DIV 11 hippocampal neurons transfected with either scrambled siRNA or GW182 siRNA at DIV7, n=22-23 neurons from 4 independent cultures, Unpaired t-test with Welch’s correction.

**Figure 5 Supplementary.**
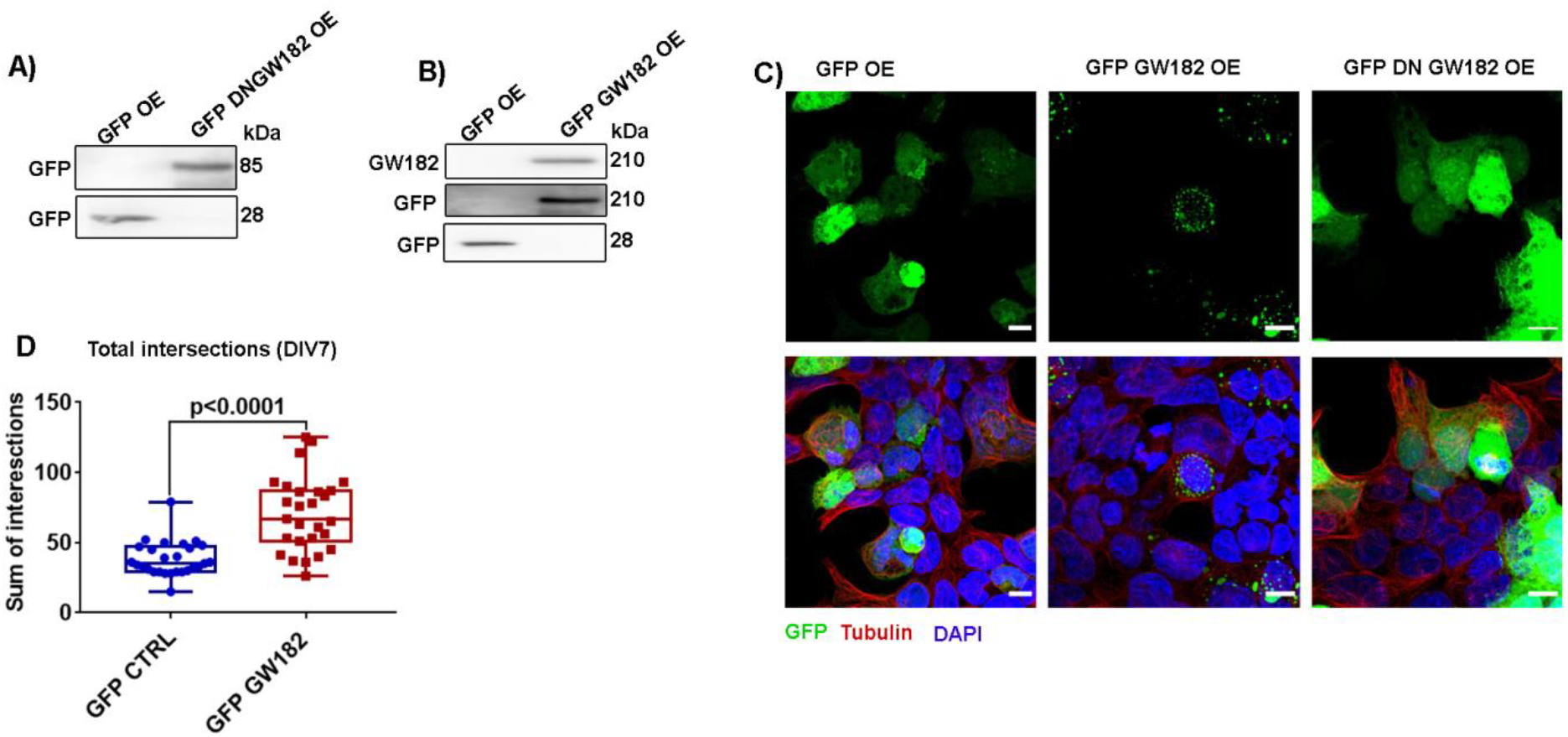
A) Immunoblot validation of GFP DNGW182 overexpression in HEK293T cells using GFP antibody. B) Immunoblot validation of GFP GW182 overexpression in HEK293T cells using GFP and GW182 antibody. C) Immunostaining characterization of GFP GW182 and GFP DNGW182 overexpression in HEK293T cells. Cells were stained with DAPI and tubulin to identify nuclear and cytosolic compartments. Scale bar represents 10 microns. D) Quantification of total intersections of DIV7 hippocampal neurons transfected with either GFP or GFP GW182 at DIV 3, n=22-23 neurons from 4 independent cultures, Unpaired t-test with Welch’s correction.

**Figure 6 Supplementary.**
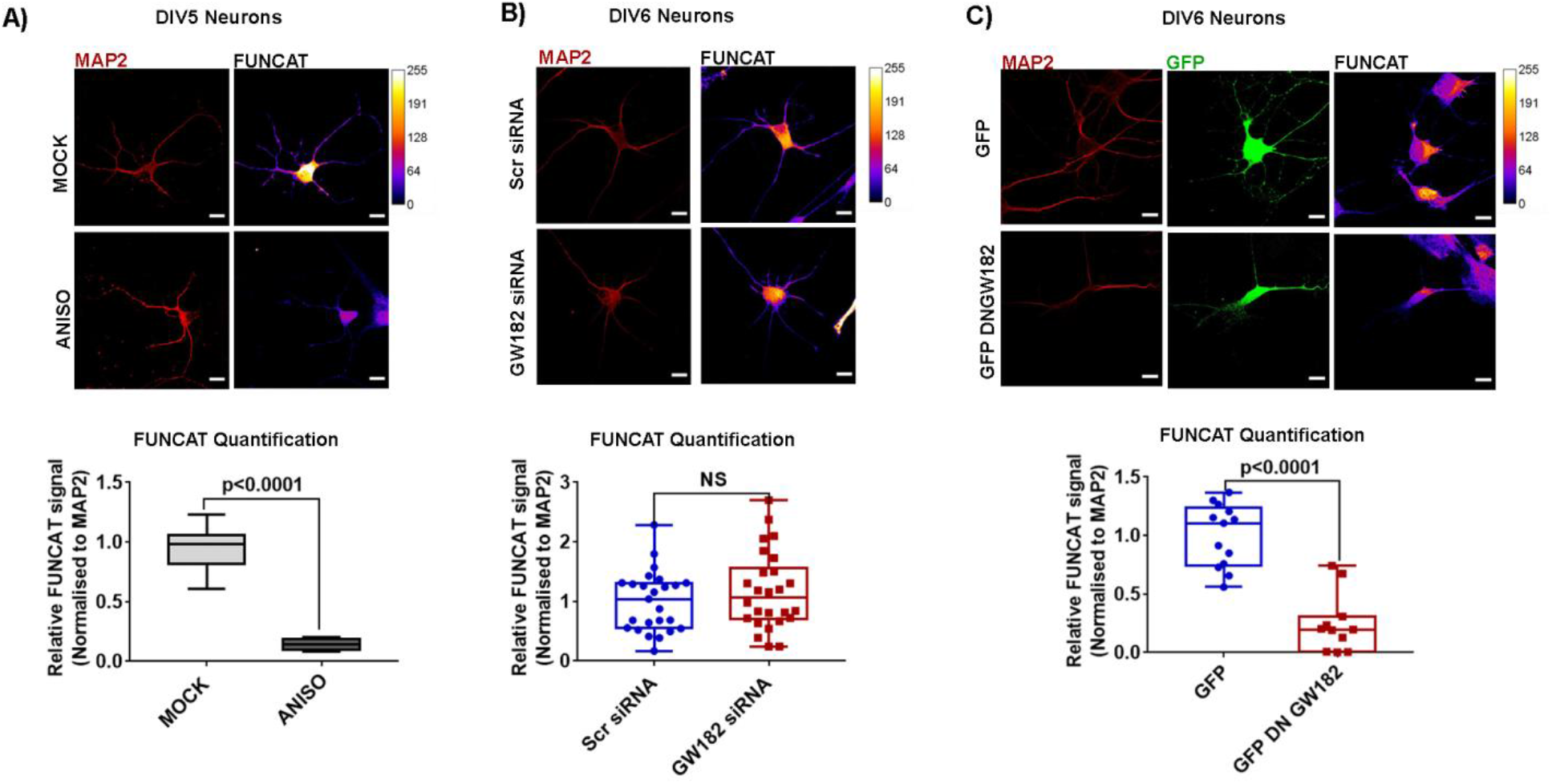
A. Representative FUNCAT images (top) and corresponding quantification of either Mock or Anisomycin treated DIV5 cultured hippocampal neurons, Scale bar represents 10 microns, n=6 neurons from 1 experiment, Unpaired t-test. B. Representative FUNCAT images (top) and corresponding quantification (bottom) of FUNCAT signal (Normalized to corresponding MAP2 signal) in DIV6 cultured hippocampal neurons, transfected with either Scrambled siRNA or GW182 siRNA on DIV3, Scale bar represents 10 microns, n=25-30 neurons from 3 independent experiments, Unpaired t-test with Welch’s correction. C. Representative FUNCAT images (top) and corresponding quantification (bottom) of FUNCAT signal (Normalized to corresponding MAP2 signal) in DIV6 cultured hippocampal neurons, transfected with either GFP control or GFP DNGW182 on DIV3, Scale bar represents 10 microns, n=25-30 neurons from 3 independent experiments, Unpaired t-test with Welch’s correction.

**Figure 7 Supplementary.**
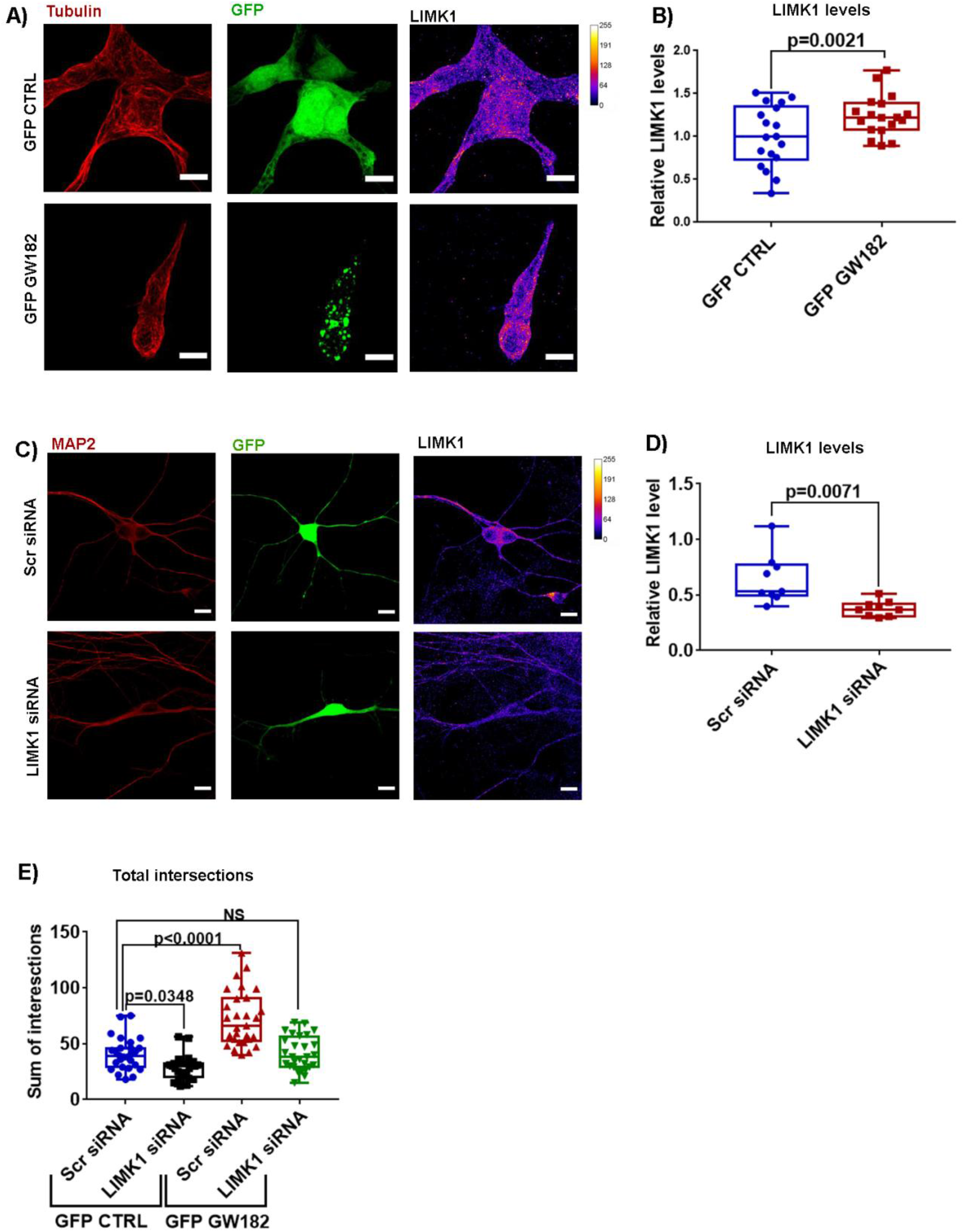
A) Representative images showing LIMK1 staining in HEK293T cells with either GFP or GFP GW182 overexpression. Scale bar represents 10 microns. B) Quantification of LIMK1 intensity in GFP or GFP GW182 overexpressing HEK293T cells. C) Immunostaining validation of LIMK1 siRNA in cultured hippocampal neurons using LIMK1 antibody. Scale bar represents 10 microns. n=18 cells, unpaired t-test. D) Quantification of LIMK1 intensity (Normalized to MAP2 intensity of corresponding neurons) in neurons transfected with either scrambled siRNA or LIMK1 siRNA. n=9 neurons from 1experiment, unpaired t-test with Welch’s correction. E) Quantification of total intersections of cultured hippocampal neurons transfected with either control GFP or GFP GW182 along with scrambled siRNA or LIMK1 siRNA. n=26-27 neurons from 3 independent experiments, One way ANOVA followed by Bonferroni’s multiple comparisons test.

## References

1. Arikkath, J., 2012. Molecular mechanisms of dendrite morphogenesis. Front. Cell. Neurosci. 6. https://doi.org/10.3389/fncel.2012.00061

2. Badhwar, A., Brown, R., Stanimirovic, D.B., Haqqani, A.S., Hamel, E., 2017. Proteomic differences in brain vessels of Alzheimer’s disease mice: Normalization by PPARγ agonist pioglitazone. J Cereb Blood Flow Metab 37, 1120–1136. https://doi.org/10.1177/0271678X16655172

3. Banerjee, S., Neveu, P., Kosik, K.S., 2009. A coordinated local translational control point at the synapse involving relief from silencing and MOV10 degradation. Neuron 64, 871–884. https://doi.org/10.1016/j.neuron.2009.11.023

4. Bartel, D.P., 2018a. Metazoan MicroRNAs. Cell 173, 20–51. https://doi.org/10.1016/j.cell.2018.03.006

5. Bartel, D.P., 2018b. Metazoan MicroRNAs. Cell 173, 20–51. https://doi.org/10.1016/j.cell.2018.03.006

6. Bateup, H.S., Denefrio, C.L., Johnson, C.A., Saulnier, J.L., Sabatini, B.L., 2013. Temporal dynamics of a homeostatic pathway controlling neural network activity. Front. Mol. Neurosci. 6. https://doi.org/10.3389/fnmol.2013.00028

7. Bestman, J.E., Cline, H.T., 2008. The RNA binding protein CPEB regulates dendrite morphogenesis and neuronal circuit assembly in vivo. Proceedings of the National Academy of Sciences 105, 20494–20499. https://doi.org/10.1073/pnas.0806296105

8. Brechbiel, J.L., Gavis, E.R., 2008. Spatial Regulation of nanos Is Required for Its Function in Dendrite Morphogenesis. Current Biology 18, 745–750. https://doi.org/10.1016/j.cub.2008.04.033

9. Chen, Y.-C., Chang, Y.-W., Huang, Y.-S., 2019. Dysregulated Translation in Neurodevelopmental Disorders: An Overview of Autism-Risk Genes Involved in Translation. Developmental Neurobiology 79, 60–74. https://doi.org/10.1002/dneu.22653

10. Chihara, T., Luginbuhl, D., Luo, L., 2007. Cytoplasmic and mitochondrial protein translation in axonal and dendritic terminal arborization. Nat Neurosci 10, 828–837. https://doi.org/10.1038/nn1910

11. Cougot, N., Bhattacharyya, S.N., Tapia-Arancibia, L., Bordonné, R., Filipowicz, W., Bertrand, E., Rage, F., 2008. Dendrites of mammalian neurons contain specialized P-body-like structures that respond to neuronal activation. J Neurosci 28, 13793–13804. https://doi.org/10.1523/JNEUROSCI.4155-08.2008

12. Crino, P.B., Eberwine, J., 1996. Molecular Characterization of the Dendritic Growth Cone: Regulated mRNA Transport and Local Protein Synthesis. Neuron 17, 1173–1187. https://doi.org/10.1016/S0896-6273(00)80248-2

13. Dieck, S. tom, Müller, A., Nehring, A., Hinz, F.I., Bartnik, I., Schuman, E.M., Dieterich, D.C., 2012. Metabolic Labeling with Noncanonical Amino Acids and Visualization by Chemoselective Fluorescent Tagging. Curr Protoc Cell Biol 0 7, Unit7.11. https://doi.org/10.1002/0471143030.cb0711s56

14. Dieterich, D.C., Hodas, J.J.L., Gouzer, G., Shadrin, I.Y., Ngo, J.T., Triller, A., Tirrell, D.A., Schuman, E.M., 2010. In situ visualization and dynamics of newly synthesized proteins in rat hippocampal neurons. Nat Neurosci 13, 897–905. https://doi.org/10.1038/nn.2580

15. Ding, L., Han, M., 2007. GW182 family proteins are crucial for microRNA-mediated gene silencing. Trends in Cell Biology 17, 411–416. https://doi.org/10.1016/j.tcb.2007.06.003

16. Dong, X., Shen, K., Bülow, H.E., 2015. Intrinsic and Extrinsic Mechanisms of Dendritic Morphogenesis. Annu. Rev. Physiol. 77, 271–300. https://doi.org/10.1146/annurev-physiol-021014-071746

17. Duchaine, T.F., Fabian, M.R., 2019. Mechanistic Insights into MicroRNA-Mediated Gene Silencing. Cold Spring Harb Perspect Biol 11. https://doi.org/10.1101/cshperspect.a032771

18. Eising, E., Carrion-Castillo, A., Vino, A., Strand, E.A., Jakielski, K.J., Scerri, T.S., Hildebrand, M.S., Webster, R., Ma, A., Mazoyer, B., Francks, C., Bahlo, M., Scheffer, I.E., Morgan, A.T., Shriberg, L.D., Fisher, S.E., 2019. A set of regulatory genes co-expressed in embryonic human brain is implicated in disrupted speech development. Molecular Psychiatry 24, 1065–1078. https://doi.org/10.1038/s41380-018-0020-x

19. Eulalio, A., Huntzinger, E., Izaurralde, E., 2008. GW182 interaction with Argonaute is essential for miRNA-mediated translational repression and mRNA decay. Nature Structural & Molecular Biology 15, 346–353. https://doi.org/10.1038/nsmb.1405

20. Eulalio, A., Tritschler, F., Izaurralde, E., 2009. The GW182 protein family in animal cells: New insights into domains required for miRNA-mediated gene silencing. RNA 15, 1433–1442. https://doi.org/10.1261/rna.1703809

21. Filipowicz, W., Bhattacharyya, S.N., Sonenberg, N., 2008. Mechanisms of post-transcriptional regulation by microRNAs: are the answers in sight? Nature Reviews Genetics 9, 102–114. https://doi.org/10.1038/nrg2290

22. Ford, L., Ling, E., Kandel, E.R., Fioriti, L., 2019. CPEB3 inhibits translation of mRNA targets by localizing them to P bodies. PNAS 116, 18078–18087. https://doi.org/10.1073/pnas.1815275116

23. Gebert, L.F.R., MacRae, I.J., 2019. Regulation of microRNA function in animals. Nature Reviews Molecular Cell Biology 20, 21–37. https://doi.org/10.1038/s41580-018-0045-7

24. Georges, P.C., Hadzimichalis, N.M., Sweet, E.S., Firestein, B.L., 2008. The Yin–Yang of Dendrite Morphology: Unity of Actin and Microtubules. Mol Neurobiol 38, 270–284. https://doi.org/10.1007/s12035-008-8046-8

25. Gorski, J.A., Zeiler, S.R., Tamowski, S., Jones, K.R., 2003. Brain-Derived Neurotrophic Factor Is Required for the Maintenance of Cortical Dendrites. J. Neurosci. 23, 6856–6865. https://doi.org/10.1523/JNEUROSCI.23-17-06856.2003

26. Guerrini, R., Mei, D., 2018. Unstable non-coding pentanucleotide repeats destabilize cortical excitability. Brain 141, 2232–2235. https://doi.org/10.1093/brain/awy196

27. Hausser, M., 2000. Diversity and Dynamics of Dendritic Signaling. Science 290, 739–744. https://doi.org/10.1126/science.290.5492.739

28. Hicks, J.A., Li, L., Matsui, M., Chu, Y., Volkov, O., Johnson, K.C., Corey, D.R., 2017. Human GW182 Paralogs are the Central Organizers for RNA-Mediated Control of Transcription. Cell Rep 20, 1543–1552. https://doi.org/10.1016/j.celrep.2017.07.058

29. Huang, Y.-W.A., Ruiz, C.R., Eyler, E.C.H., Lin, K., Meffert, M.K., 2012. Dual Regulation of miRNA Biogenesis Generates Target Specificity in Neurotrophin-Induced Protein Synthesis. Cell 148, 933–946. https://doi.org/10.1016/j.cell.2012.01.036

30. Huntzinger, E., Kuzuoğlu-Öztürk, D., Braun, J.E., Eulalio, A., Wohlbold, L., Izaurralde, E., 2013. The interactions of GW182 proteins with PABP and deadenylases are required for both translational repression and degradation of miRNA targets.Nucleic Acids Res 41, 978–994. https://doi.org/10.1093/nar/gks1078

31. Jakymiw, A., Lian, S., Eystathioy, T., Li, S., Satoh, M., Hamel, J.C., Fritzler, M.J., Chan, E.K.L., 2005. Disruption of GW bodies impairs mammalian RNA interference. Nature Cell Biology 7, 1267–1274. https://doi.org/10.1038/ncb1334

32. Jan, Y.-N., Jan, L.Y., 2010. Branching out: mechanisms of dendritic arborization. Nat Rev Neurosci 11, 316–328. https://doi.org/10.1038/nrn2836

33. Jaworski, J., Spangler, S., Seeburg, D.P., Hoogenraad, C.C., Sheng, M., 2005. Control of Dendritic Arborization by the Phosphoinositide-3’-Kinase–Akt–Mammalian Target of Rapamycin Pathway. J Neurosci 25, 11300–11312. https://doi.org/10.1523/JNEUROSCI.2270-05.2005

34. Kaech, S., Banker, G., 2006. Culturing hippocampal neurons. Nature Protocols 1, 2406–2415. https://doi.org/10.1038/nprot.2006.356

35. Keil, K.P., Miller, G.W., Chen, H., Sethi, S., Schmuck, M.R., Dhakal, K., Kim, J.W., Lein, P.J., 2018. PCB 95 promotes dendritic growth in primary rat hippocampal neurons via mTOR-dependent mechanisms. Arch Toxicol 92, 3163–3173. https://doi.org/10.1007/s00204-018-2285-x

36. Kenny, P.J., Zhou, H., Kim, M., Skariah, G., Khetani, R.S., Drnevich, J., Arcila, M.L., Kosik, K.S., Ceman, S., 2014. MOV10 and FMRP regulate AGO2 association with microRNA recognition elements. Cell Rep 9, 1729–1741. https://doi.org/10.1016/j.celrep.2014.10.054

37. Konietzny, A., Bär, J., Mikhaylova, M., 2017. Dendritic Actin Cytoskeleton: Structure, Functions, and Regulations. Front Cell Neurosci 11. https://doi.org/10.3389/fncel.2017.00147

38. Kosik, K.S., 2006. The neuronal microRNA system. Nature Reviews Neuroscience 7, 911–920. https://doi.org/10.1038/nrn2037

39. Kulkarni, V.A., Firestein, B.L., 2012. The dendritic tree and brain disorders. Molecular and Cellular Neuroscience 11.

40. Kumar, V., Zhang, M.-X., Swank, M.W., Kunz, J., Wu, G.-Y., 2005. Regulation of Dendritic Morphogenesis by Ras–PI3K–Akt–mTOR and Ras–MAPK Signaling Pathways. J. Neurosci. 25, 11288–11299. https://doi.org/10.1523/JNEUROSCI.2284-05.2005

41. Kute, P.M., Ramakrishna, S., Neelagandan, N., Chattarji, S., Muddashetty, R.S., 2019. NMDAR mediated translation at the synapse is regulated by MOV10 and FMRP. Mol Brain 12, 65. https://doi.org/10.1186/s13041-019-0473-0

42. La Rocca, G., Olejniczak, S.H., González, A.J., Briskin, D., Vidigal, J.A., Spraggon, L., DeMatteo, R.G., Radler, M.R., Lindsten, T., Ventura, A., Tuschl, T., Leslie, C.S., Thompson, C.B., 2015. In vivo, Argonaute-bound microRNAs exist predominantly in a reservoir of low molecular weight complexes not associated with mRNA. Proc Natl Acad Sci USA 112, 767–772. https://doi.org/10.1073/pnas.1424217112

43. Lee, A., Li, W., Xu, K., Bogert, B.A., Su, K., Gao, F.-B., 2003. Control of dendritic development by the Drosophila fragile X-related gene involves the small GTPase Rac1. Development 130, 5543–5552. https://doi.org/10.1242/dev.00792

44. Lein, P.J., Higgins, D., 1991. Protein synthesis is required for the initiation of dendritic growth in embryonic rat sympathetic neurons in vitro. Developmental Brain Research 60, 187–196. https://doi.org/10.1016/0165-3806(91)90047-M

45. Li, S., Wang, L., Fu, B., Berman, M.A., Diallo, A., Dorf, M.E., 2014. TRIM65 regulates microRNA activity by ubiquitination of TNRC6. Proc Natl Acad Sci U S A 111, 6970–6975. https://doi.org/10.1073/pnas.1322545111

46. Liu, J., Rivas, F.V., Wohlschlegel, J., Yates, J.R., Parker, R., Hannon, G.J., 2005. A role for the P-body component, GW182, in microRNA function. Nat Cell Biol 7, 1261–1266. https://doi.org/10.1038/ncb1333

47. Luo, Y., Na, Z., Slavoff, S.A., 2018. P-Bodies: Composition, Properties, and Functions. Biochemistry 57, 2424–2431. https://doi.org/10.1021/acs.biochem.7b01162

48. Martínez-Cerdeño, V., 2017. Dendrite and spine modifications in autism and related neurodevelopmental disorders in patients and animal models: Dendrite and Spine in Autism. Devel Neurobio 77, 393–404. https://doi.org/10.1002/dneu.22417

49. McAllister, A.K., Katz, L.C., Lo, D.C., 1996. Neurotrophin Regulation of Cortical Dendritic Growth Requires Activity. Neuron 17, 1057–1064. https://doi.org/10.1016/S0896-6273(00)80239-1

50. McNeill, E., Van Vactor, D., 2012. microRNAs Shape the Neuronal Landscape. Neuron 75, 363–379. https://doi.org/10.1016/j.neuron.2012.07.005

51. Meijering, E., Jacob, M., Sarria, J.-C.F., Steiner, P., Hirling, H., Unser, M., 2004. Design and validation of a tool for neurite tracing and analysis in fluorescence microscopy images. Cytometry Part A 58A, 167–176. https://doi.org/10.1002/cyto.a.20022

52. Muddashetty, R.S., Nalavadi, V.C., Gross, C., Yao, X., Xing, L., Laur, O., Warren, S.T., Bassell, G.J., 2011. Reversible inhibition of PSD-95 mRNA translation by miR-125a, FMRP phosphorylation and mGluR signaling. Mol Cell 42, 673–688. https://doi.org/10.1016/j.molcel.2011.05.006

53. Nawalpuri, B., Ravindran, S., Muddashetty, R.S., 2020. The Role of Dynamic miRISC During Neuronal Development. Front Mol Biosci 7. https://doi.org/10.3389/fmolb.2020.00008

54. Niaz, S., Hussain, M.U., 2018. Role of GW182 protein in the cell. The International Journal of Biochemistry & Cell Biology 101, 29–38. https://doi.org/10.1016/j.biocel.2018.05.009

55. Olejniczak, S.H., Rocca, G.L., Radler, M.R., Egan, S.M., Xiang, Q., Garippa, R., Thompson, C.B., 2016. Coordinated Regulation of Cap-Dependent Translation and MicroRNA Function by Convergent Signaling Pathways. Molecular and Cellular Biology 36, 2360–2373. https://doi.org/10.1128/MCB.01011-15

56. Perycz, M., Urbanska, A.S., Krawczyk, P.S., Parobczak, K., Jaworski, J., 2011. Zipcode Binding Protein 1 Regulates the Development of Dendritic Arbors in Hippocampal Neurons. J. Neurosci. 31, 5271–5285. https://doi.org/10.1523/JNEUROSCI.2387-10.2011

57. Pfaff, J., Hennig, J., Herzog, F., Aebersold, R., Sattler, M., Niessing, D., Meister, G., 2013. Structural features of Argonaute–GW182 protein interactions. PNAS 110, E3770–E3779. https://doi.org/10.1073/pnas.1308510110

58. Poulain, F.E., Sobel, A., 2010. The microtubule network and neuronal morphogenesis: Dynamic and coordinated orchestration through multiple players. Molecular and Cellular Neuroscience 43, 15–32. https://doi.org/10.1016/j.mcn.2009.07.012

59. Rajgor, D., Sanderson, T.M., Amici, M., Collingridge, G.L., Hanley, J.G., 2018. NMDAR-dependent Argonaute 2 phosphorylation regulates miRNA activity and dendritic spine plasticity. EMBO J 37. https://doi.org/10.15252/embj.201797943

60. Rajman, M., Schratt, G., 2017. MicroRNAs in neural development: from master regulators to fine-tuners. Development 144, 2310–2322. https://doi.org/10.1242/dev.144337

61. Ramakrishna, S., Muddashetty, R.S., 2019. Emerging Role of microRNAs in Dementia. Journal of Molecular Biology, Dementia, Brain Disorders and Molecular Mechanisms 431, 1743–1762. https://doi.org/10.1016/j.jmb.2019.01.046

62. Ravindran, S., Nalavadi, V.C., Muddashetty, R.S., 2019. BDNF Induced Translation of Limk1 in Developing Neurons Regulates Dendrite Growth by Fine-Tuning Cofilin1 Activity. Front. Mol. Neurosci. 12. https://doi.org/10.3389/fnmol.2019.00064

63. Saito, A., Miyajima, K., Akatsuka, J., Kondo, H., Mashiko, T., Kiuchi, T., Ohashi, K., Mizuno, K., 2013. CaMKIIβ-mediated LIM-kinase activation plays a crucial role in BDNF-induced neuritogenesis. Genes to Cells 18, 533–543. https://doi.org/10.1111/gtc.12054

64. Schratt, G.M., Tuebing, F., Nigh, E.A., Kane, C.G., Sabatini, M.E., Kiebler, M., Greenberg, M.E., 2006. A brain-specific microRNA regulates dendritic spine development. Nature 439, 283–289. https://doi.org/10.1038/nature04367

65. Skalecka, A., Liszewska, E., Bilinski, R., Gkogkas, C., Khoutorsky, A., Malik, A.R., Sonenberg, N., Jaworski, J., 2016. mTOR kinase is needed for the development and stabilization of dendritic arbors in newly born olfactory bulb neurons. Developmental Neurobiology 76, 1308–1327. https://doi.org/10.1002/dneu.22392

66. Skariah, G., Seimetz, J., Norsworthy, M., Lannom, M.C., Kenny, P.J., Elrakhawy, M., Forsthoefel, C., Drnevich, J., Kalsotra, A., Ceman, S., 2017. Mov10 suppresses retroelements and regulates neuronal development and function in the developing brain. BMC Biology 15, 54. https://doi.org/10.1186/s12915-017-0387-1

67. Slomnicki, L.P., Pietrzak, M., Vashishta, A., Jones, J., Lynch, N., Elliot, S., Poulos, E., Malicote, D., Morris, B.E., Hallgren, J., Hetman, M., 2016. Requirement of Neuronal Ribosome Synthesis for Growth and Maintenance of the Dendritic Tree* 291, 20.

68. Sternburg, E.L., Estep, J.A., Nguyen, D.K., Li, Y., Karginov, F.V., 2018. Antagonistic and cooperative AGO2-PUM interactions in regulating mRNAs. Scientific Reports 8, 15316. https://doi.org/10.1038/s41598-018-33596-4

69. Tiedge, H., Brosius, J., 1996. Translational Machinery in Dendrites of Hippocampal Neurons in Culture. J. Neurosci. 16, 7171–7181. https://doi.org/10.1523/JNEUROSCI.16-22-07171.1996

70. Wolterhoff, N., Gigengack, U., Rumpf, S., 2020. PP2A phosphatase is required for dendrite pruning via actin regulation in Drosophila. EMBO reports 21, e48870. https://doi.org/10.15252/embr.201948870

71. Xing, L., Yao, X., Williams, K.R., Bassell, G.J., 2012. Negative regulation of RhoA translation and signaling by hnRNP-Q1 affects cellular morphogenesis. Molecular Biology of the Cell 23, 10.

72. Xu, Jing, Du, Y., Xu, Jing-wei, Hu, X., Gu, L., Li, X., Hu, P., Liao, T., Xia, Q., Sun, Q., Shi, L., Luo, J., Xia, J., Wang, Z., Xu, Junyu, 2019. Neuroligin 3 Regulates Dendritic Outgrowth by Modulating Akt/mTOR Signaling. Front. Cell. Neurosci. 13. https://doi.org/10.3389/fncel.2019.00518

73. Yao, B., Li, S., Jung, H.M., Lian, S.L., Abadal, G.X., Han, F., Fritzler, M.J., Chan, E.K.L., 2011. Divergent GW182 functional domains in the regulation of translational silencing. Nucleic Acids Res 39, 2534–2547. https://doi.org/10.1093/nar/gkq1099

74. Ye, B., Petritsch, C., Clark, I.E., Gavis, E.R., Jan, L.Y., Jan, Y.N., 2004. nanos and pumilio Are Essential for Dendrite Morphogenesis in Drosophila Peripheral Neurons. Current Biology 14, 314–321. https://doi.org/10.1016/j.cub.2004.01.052

75. Zekri, L., Huntzinger, E., Heimstädt, S., Izaurralde, E., 2009. The Silencing Domain of GW182 Interacts with PABPC1 To Promote Translational Repression and Degradation of MicroRNA Targets and Is Required for Target Release. Molecular and Cellular Biology 29, 6220–6231. https://doi.org/10.1128/MCB.01081-09

76. Zielezinski, A., Karlowski, W.M., 2015. Early origin and adaptive evolution of the GW182 protein family, the key component of RNA silencing in animals. RNA Biol 12, 761–770. https://doi.org/10.1080/15476286.2015.1051302

